# New perspectives on the calculation of bioaccumulation metrics for active substances in living organisms

**DOI:** 10.1101/2020.07.07.185835

**Authors:** Aude Ratier, Christelle Lopes, Gauthier Multari, Vanessa Mazerolles, Patrice Carpentier, Sandrine Charles

## Abstract

Today, only few ready-to-use and convenient decision-making tools are available in ecotoxicology concerning accumulation and effects of chemical substances on organisms, accounting for exposure situations that are known to be complex (routes of exposure, metabolism, mixtures, etc.). This paper presents new perspectives on the generic calculation of bioaccumulation metrics via the innovative web tool MOSAIC_bioacc_ (http://mosaic.univ-lyon1.fr/bioacc). MOSAIC_bioacc_ provides all kind of bioaccumulation metrics associated with their uncertainty whatever the species-compound combination. MOSAIC_bioacc_ expects accumulation-depuration data as inputs, even with complex exposure and clearance patterns, to quickly perform their relevant analysis. MOSAIC_bioacc_ intends to facilitate the daily work of regulators, or any ecotoxicologist, who will freely benefit from a user-friendly on-line interface that automatically fits toxicokinetic models without needs for users to invest in the technical aspects to get bioaccumulation metrics estimates. MOSAIC_bioacc_ also provides all results in a fully transparent way to ensure reproducibility.

## INTRODUCTION

Faced with the current environmental challenges, linked in particular to environmental pollution, ecotoxicology must today provide relevant and effective decision-making tools regarding bioaccumulation and effects of chemical substances on living organisms. Such tools must account for various exposure situations, environmentally realistic but complex (*e.g*., several routes of exposure, metabolism of substances, mixtures, etc.). Among available methods, toxicokinetic-toxicodynamic (TKTD) models are now strongly recommended to describe the link between exposure concentrations and effects on individual life-history traits over time from experimental data collected through toxicity tests, even standard ones (EFSA PPR Panel 2018). More specifically, the TK part of these models is used to relate the exposure concentration to the time course of the internal concentration within organisms, considering various processes such as accumulation, depuration, metabolization and excretion (known as ADME processes). As some recent regulations, the EU regulation No 283/2013 for plant protection products in marketing authorisation applications requires for example a bioaccumulation test on fish according to OECD Test guideline 305 (OECD 2012), which consists in an accumulation phase followed by a depuration phase. During the accumulation phase, fish are exposed to a substance of interest at a range of concentrations, chosen according to the assumed mode of action of the substance. After a certain time period fixed by the experimenter, organisms are transferred to a clean medium for a depuration phase. The concentration of the substance (and of its potential metabolites) is followed within fish at regular time points during both phases leading *in fine* to the estimation of bioaccumulation metrics. In this paper, for the sake of generality, we chose the generic expression “bioaccumulation metrics” to denote either bioconcentration factors (BCF) used when exposure is via water, biota-sediment accumulation factors (BSAF) when exposure is via sediment or biomagnification factors (BMF) when exposure is via food. Bioaccumulation metrics appeared to us as the best compromise regarding the wide diversity of terms used in the scientific literature (for example, USEPA (1994), Gobas et al. (2009) and Burkhard et al. (2012)). Nevertheless, preference is often given to experimentally derived BCF estimates to be used for secondary poisoning assessment under Biocidal Products Regulation (European Commission 2012). Bioaccumulation tests are of course not only limited to fish, even if a test according to OECD Test guideline 305 (OECD 2012) is preferred when experimental information on bioaccumulation is needed for PBT/vPvB assessment under REACH regulation (ECHA 2017; European Commission 2006). Consequently, any other species can be used, such as benthic invertebrates, terrestrial oligochaetes or birds, depending on the substance under consideration. From a regulatory point of view, bioaccumulation metrics are key decision criteria used to evaluate concentrations of active substances in food items of vertebrates (especially piscivorous birds and mammals), making the estimation of these metrics with the most precision as possible a highly crucial methodological challenge.

All bioaccumulation metrics rely on estimates of kinetic parameters as involved in toxicokinetic (TK) models. In the past decades, many types of methods have been proposed to get these estimates from simple TK models, most of them providing BCF estimates separately considering the kinetics for both accumulation and depuration phases as observed in dedicated experiments (OECD 2012). Nevertheless, TK model parameters are known to be highly correlated, so that separating their estimation prevents to account for a mutual influence on their uncertainty. Moreover, kinetic parameter estimates are only rarely provided with their uncertainty, although this is now expected by the regulatory bodies (EFSA Scientific Committee 2018). Consequently, concomitantly to the above-mentioned challenge, environmental risk assessment could be improved if complete tools allowing for a simultaneously estimation of all TK model parameters associated with their uncertainty would be available in support of stakeholders that need to fulfil regulatory expectations. These tools must also be easy to use to overcome the scepticism of regulators who are faced with multiple TK models and implementation methods, while thinking about their standardization at the same time (Tan et al. 2020). An R-code was first proposed in 2016 (Aldenberg 2019) that allowed to analyse data collected only from the OCED test guideline 305. In the same line of thought, a spreadsheet was recently proposed by Gobas et al. (2020). Our paper goes beyond by considering all type of species-compound accumulation-depuration data (not only those of the OECD test guideline 305) which analysis leads to one or several bioaccumulation metrics of interest.

Ratier et al. (2019) recently proposed a full revisit of the TK modelling approach based on a unified inference method to estimate parameters of TK models for both accumulation and depuration phases, simultaneously, automatically associating the uncertainties. This innovative framework has been thought to make it possible to further incorporate the TK part into complete TKTD models. Benefiting from this innovation, we present new perspectives for a facilitated calculation of any type of bioaccumulation metrics (such as BCF/BSAF/BMF) thanks to the new ready-to-use statistical web tool MOSAIC_bioacc_ (http://mosaic.univ-lyon1.fr/bioacc). MOSAIC_bioacc_ runs one-compartment TK models that are automatically designed according to the input data. MOSAIC_bioacc_ leads to bioaccumulation metrics associated with their uncertainty propagated from the kinetic parameter estimates, without the need for users to invest underlying technical aspects. MOSAIC_bioacc_ is free of use, fully integrated within the all-in-one facility MOSAIC itself (http://mosaic.univ-lyon1.fr). MOSAIC_bioacc_ is regularly updated to always offers the very latest conceptual advances related to TK models. Today, MOSAIC_bioacc_ allows accounting for several exposure routes (water, pore water, sediment and food), for metabolism of chemicals (if the input experimental data include measurements for both the parent chemical and its metabolites), and for potential growth of organisms (if growth measurements are included within the data set). The use of MOSAIC_bioacc_ only requires users to upload their experimental data, collected via standard protocols or from home-made experimental designs. Priority was first given to the calculation of BCF/BSAF/BMF because they are the widely used bioaccumulation metrics in the current regulatory guidelines. MOSAIC_bioacc_ get their estimate as probability distributions that are summarized for users by the median (50^th^ centile of the distribution) and the 95% credible interval (delimited by the 2.5^th^ and 97.5^th^ centiles) quantifying the uncertainty. In addition, the fitting plots and all model parameter estimates are provided, followed by a collection of goodness-of-fit criteria allowing users to check the relevance of the results. All outputs can be downloaded under different formats for further inclusion into any home-made document (in particular the open-source programming code), and a full report can also be directly downloaded gathering everything that is displayed to users on the web page. These two latter features of MOSAIC_bioacc_ guarantee both reproducibility and transparency of underlying calculations.

## MATERIALS AND METHODS

MOSAIC_bioacc_ is part of the web platform MOSAIC (https://mosaic.univ-lyon1.fr/, Charles et al. 2018). It was developed as a Shiny environment (Chang et al. 2020), available at https://mosaic.univ-lyon1.fr/bioacc and hosted at the Rhône-Alpes Bioinformatics Center PRABI (PRABI 2020). A user guide and an explanatory video are immediately available in the introductory section of the application to fully assist users step by step in appropriating the tool and its features. Details on underlying ordinary differential equations and their solving are also provided within a detailed user guide (see supplemental data, **Annex 1**), thus ensuring all required transparency as recommended by EFSA for a good modelling practice (EFSA PPR Panel 2014).

### Data uploading

When using MOSAIC_bioacc_, the first step is to upload input data (**Fig. 1-a**). MOSAIC_bioacc_ expects to receive experimental exposure time-course data, including at least an accumulation phase, as a .txt file or a .csv file (comma, semicolon or tabular separator) with a specific format. Each line of the table corresponds to one time point for a given replicate and a given exposure concentration of the contaminant. The data table must contain at least four columns, with the exact following headers, the order of columns being not important (**Table 1**): ‘time’ (the time point of the measurement at the exposure concentration, in hours, minutes, days or weeks); ‘expw’, ‘exppw’, ‘exps’ or ‘expf’: the exposure concentration of the contaminant in the medium, that is water, pore water, sediment or food, respectively, all expressed in μg.mL^-1^ or in μg.g^-1^); ‘replicate’ (a number or a character that is unique for each replicate, dimensionless); and ‘conc’ (the concentration of the contaminant, and of its potential metabolites, measured within organisms, in μg.g^-1^). According to the experimental design, further columns can be added in the data file: ‘expw’, ‘exppw’, ‘exps’ and/or ‘expf’ if several exposure routes are considered together, ‘concm*ℓ*’ (the concentrations of metabolite *ℓ* derived from the parent compound within the organisms, *e.g*., concm1, concm2,…), and ‘growth’ (if growth measurements of the organisms are available).

**Figure 1.**
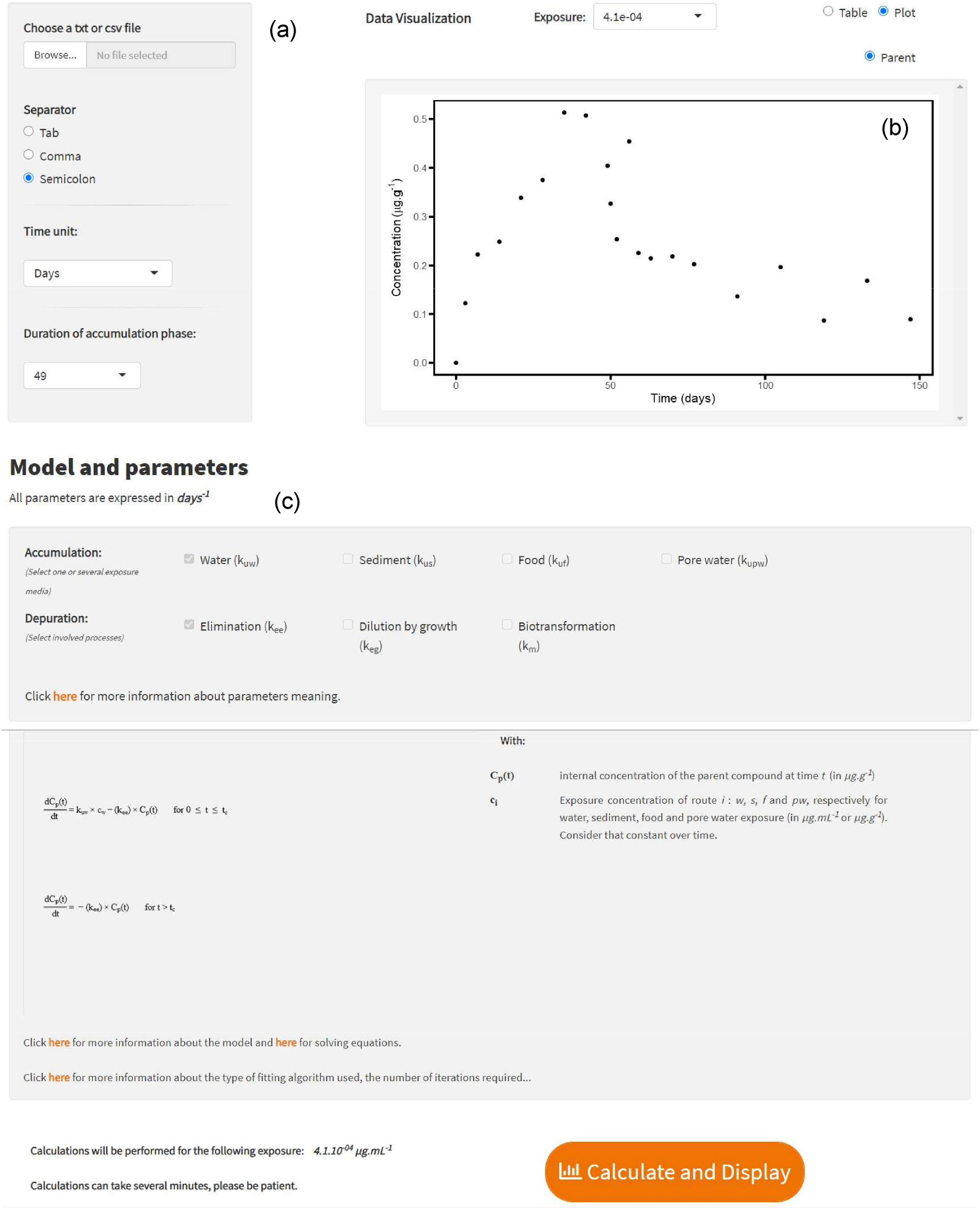
Data uploading and user information as required from the MOSAIC_bioacc_ homepage.

**Table 1.**
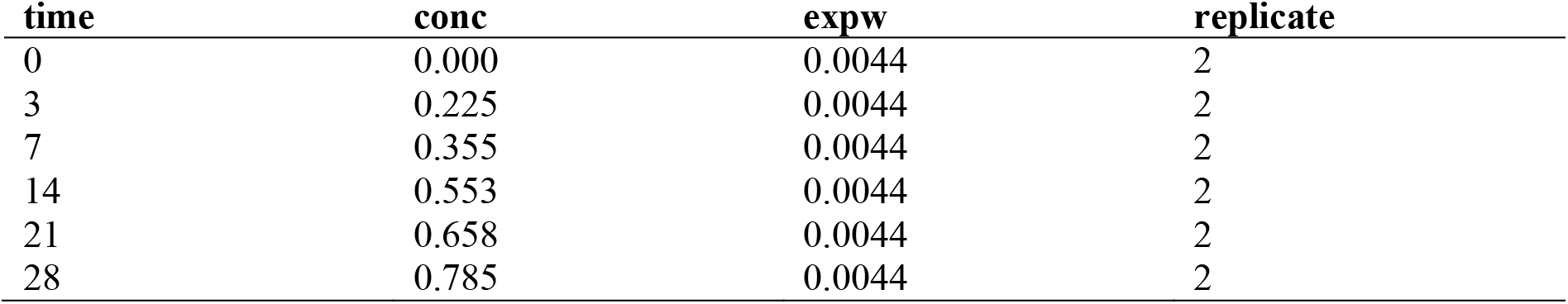
Example of a data set ready to be used in MOSAIC_bioacc_. The data set must contain four columns whatever their order: ‘time’ (time points of the measurements at the exposure concentration, in time unit: here in days), ‘conc’ (contaminant concentrations measured within organisms, that must be expressed in μg.g^-1^), ‘expw’ (the contaminant concentration in the exposure medium, here water, that must be expressed in μg.mL^-1^), and ‘replicate’ (a number or a character that is unique for each replicate, dimensionless).

As shown on **Fig. 1-a**, MOSAIC_bioacc_ users can either upload their own data set with a click on ‘Browse’ (by taking care about the expected format specification) or try MOSAIC_bioacc_ with example data sets (in total six data sets are proposed, each with different characteristics). When the upload is complete, users must manually select the appropriate separator, the time unit and the duration of the accumulation phase; please note that, when using example data sets, these fields are automatically filled in.

MOSAIC_bioacc_ first provides a table with the raw data allowing users to check if the data were correctly entered. Users can also visualize a plot with the raw data (**Fig. 1-b**). In case data were collected for several exposure concentrations and if the users have uploaded all, one must be chosen for the MOSAIC_bioacc_ analysis. Note that only one file at a time can be analysed. Also, when another exposure concentration from the same data file is chosen, the duration of the accumulation phase is reset and framed in orange to invite users for update before to launch new calculations. Example files provided in MOSAIC_bioacc_ are dedicated to assist users in formatting their own data and to appropriate the different MOSAIC_bioacc_ features from the very first step. This paper illustrates this step-by-step process based on a typical data set of a toxicokinetic bioassay where internal concentrations were collected in fathead minnows (*Pimephales promelas*) exposed via contaminated water to a highly hydrophobic chemical (logK_ow_ = 9.06) at an exposure concentration of 0.0044 μg.mL^-1^ over 49 days, with one replicate at each time point. After 49 days, minnows were transferred in a clean medium for 98 days more (Crookes and Brooke 2011). The data set (CSV format) and the report with all results (HTML format) can be downloaded directly from MOSAIC_bioacc_.

### Model and parameters

All TK models considered in MOSAIC_bioacc_ describe organisms as single compartments for which a first-order kinetic bioaccumulation model accounting for several exposure routes and elimination processes can be expressed in a generic way as follows (Eqs. (1) to (4)):

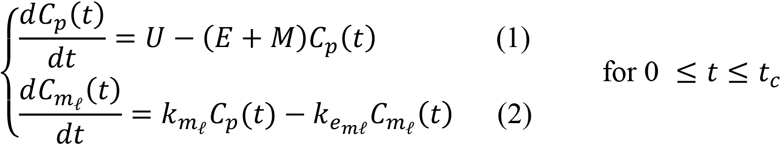

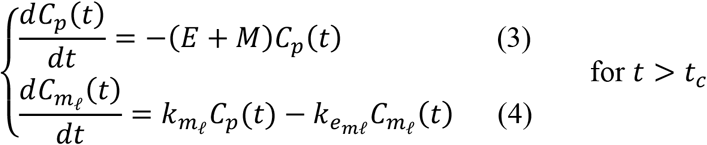

where *C_p_*(*t*) is the internal concentration of the parent compound at time *t* (μg.g^-1^), *C_mℓ_*(*t*) the internal concentration (μg.g^-1^) of metabolites (∀*ℓ* = 1…*L* with *L* the total number of metabolites) at time *t*, *U* the sum of all uptake terms, *E* the sum of all elimination terms for the parent compound, *M* the sum of all metabolization terms, *k_mℓ_* the metabolization rate of metabolite *ℓ* (*time*^-1^) and *k_e_mℓ__* the elimination rate of metabolite *ℓ* (*time*^-1^). **Table 2** gives an overview of all parameter and variable meaning. The dynamical system in equations (1) to (4), corresponding to the deterministic part of the model, can explicitly be solved when the exposure concentration is assumed to be constant over time (**Annex 1**). A Gaussian probability distribution was assumed as the stochastic part of the final model, based on the quantitative continuous nature of the concentration variables for both the parent compound and its metabolites within the organisms (Eqs. (5) and (6)):

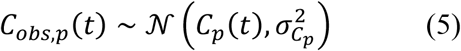

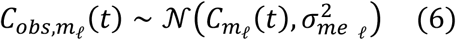

**Table 2.**
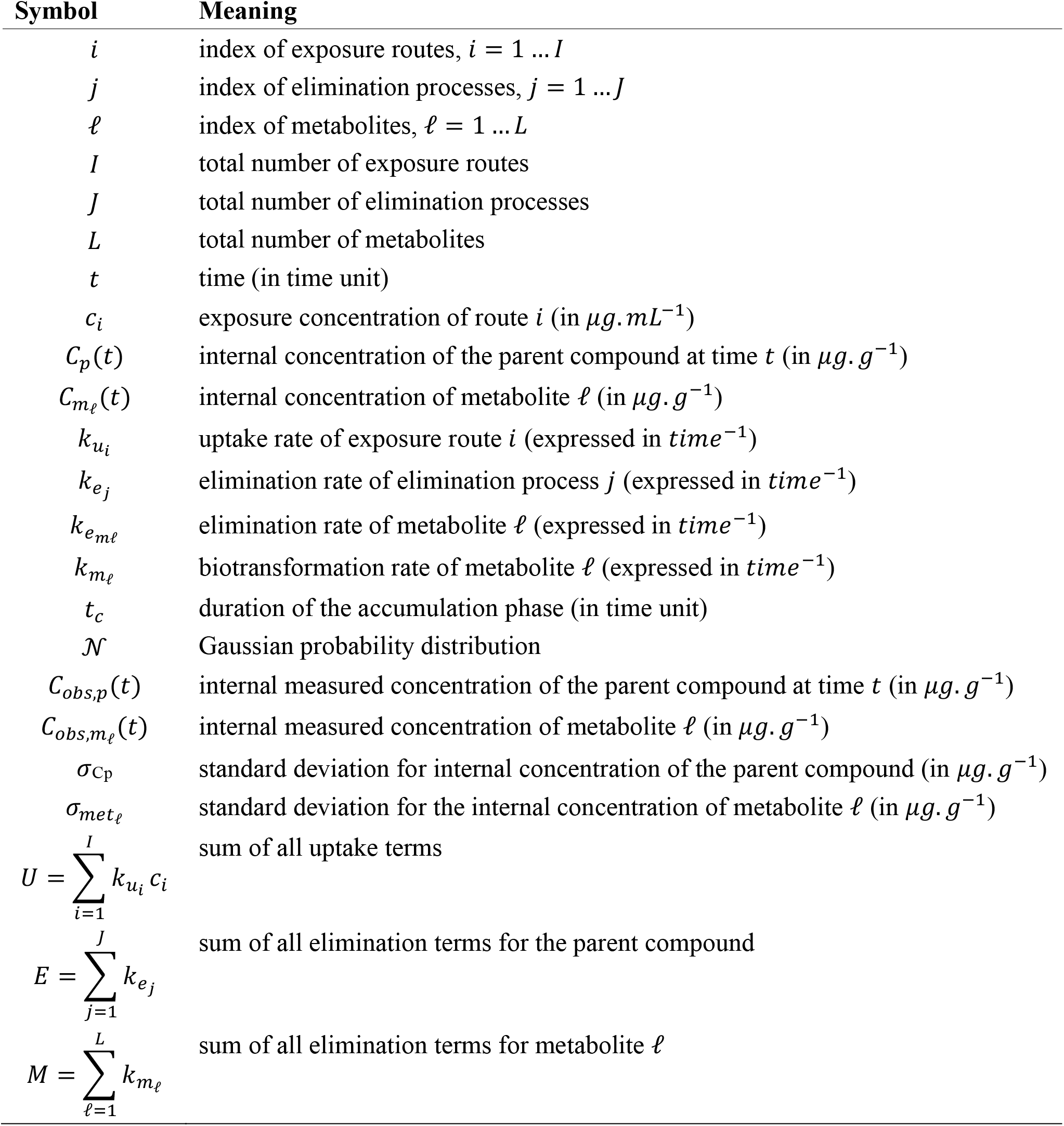
Meaning of parameters and variables of the TK model used in MOSAIC_bioacc_.

Today MOSAIC_bioacc_ proposes data analyses by including until four exposure routes via water, pore water, sediment, and/or food), until three processes of elimination, which are excretion, growth dilution and biotransformation, with a maximum of 15 metabolites directly deriving from the parent compound (*i.e*., phase I metabolism). For example, if three metabolites are considered (*L* = 3), the number of parameters involved in the most complete TK model equals 19. In total, users have 112 possible models, automatically designed according to their data (see supplemental data, **Annex 2**). For a given data set, the most complete model is built up by default and first proposed to users, from which they can perform the MOSAIC_bioacc_ analysis (**Fig. 1-c**). Users can also deselect some of the parameters (based on biological hypotheses related to the most probable exposure route or by neglecting one elimination process, for example). These choices lead to the automatic building of a nested TK sub-model to fit again on the data.

For clarity reasons, this paper illustrates MOSAIC_bioacc_ features from a simple data set (Pimephales_two.csv file) considering only the water exposure route (parameter *k_uw_* for the uptake rate) and the excretion process (parameter *k_ee_* for the elimination rate), that is one of the simplest TK models among the 112 possibilities. The corresponding equations for the deterministic part of this TK model are given below (Eqs. (7) and (8)):

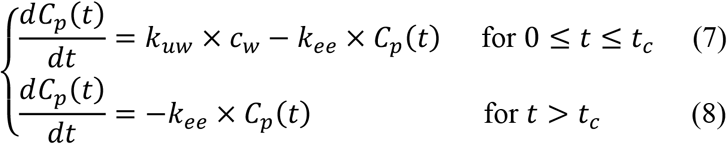

Finally, there are only three parameters to estimate: *k_uw_*, *k_ee_* and parameter σ_C_p__ of the Gaussian distribution (Eq. (5)). An illustration of a more complex data set with biotransformation and growth processes for a benthic invertebrate is given as supplemental material (**Annex 3**), as well as examples for exposure route by sediment (**Annex 4**) or food (**Annex 5**).

Once the Pimephales_two.csv example file has been uploaded, all required fields related to the experimental design are automatically filled in and raw data can be visualized, either as a table (default) or a plot, and the button to launch calculations (‘Calculate and Display’) is unlocked (**Fig. 1-c**). Once this button clicked, calculations start running with a progress bar informing users about progress. When calculations are finished, results are displayed, as plots or tables first for the bioaccumulation metrics, then for the fitting plots and some relevant goodness-of-fit criteria.

### Calculations

#### Bayesian inference

Computations underlying MOSAIC_bioacc_ results are performed with JAGS (Plummer 2019) and the R software (R Core Team 2020, version 4.0.2) via the `rjags` and `jagsUI` packages (Plummer 2019; Kellner 2019). Models are fitted to bioaccumulation data using Bayesian inference via Monte Carlo Markov Chain (MCMC) sampling. For each model, calculation running starts with a short sampling on three MCMC chains (5,000 iterations after a burn-in phase of 10,000 iterations) using the Raftery and Lewis method (Raftery and Lewis 1992) to set the necessary thinning and the appropriate number of iterations in order to reach a precise and accurate estimation of each model parameter. Thanks to `rjags`, model parameters are retrieved as a joint posterior distribution from the likelihood of the observed data combined with prior information on parameters. All details on this approach can be found in the original research paper (Ratier et al. 2019) but also in many other papers in the field of ecotoxicology (Billoir et al. 2011; EFSA PPR Panel 2018).

#### Choice of prior distributions

For simplicity reasons, we hid the choice of priors to MOSAIC_bioacc_ users; hence, they cannot be changed, except by downloading the open source programming code and handling it directly within the R software. To ensure genericity of priors we chose non-informative (−5, 5) log10-uniform distributions for all uptake and elimination rate constants, and non-informative (0, A) uniform distributions for all standard deviations with a large A, here defined as five times the maximum internal measured concentration, which is then removed from the data set, as usually proceeded (Gelman 2006).

#### Bioaccumulation metrics

Bioaccumulation metrics are the first outputs delivered by MOSAIC_bioacc_. From the example chosen for this paper, the data analysis led to both the kinetic bioconcentration factor (*BCF_k_*) and the steady state bioconcentration factor (*BCF_ss_*), with the following exact mathematical expressions (Eqs. (9) and (10)):

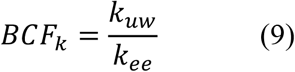

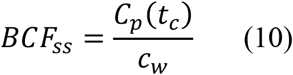

where *C_p_*(*t_c_*) is the internal parent compound concentration (in μg.g^-1^) at the end of the accumulation phase (that is at *t* = *t_c_*, in time) and *c_w_* is the exposure contaminant concentration in water (μg.mL^-1^). More details about calculations of bioaccumulation metrics at steady state are provided in **Annex 6**.

## RESULTS AND DISCUSSION

### Bioaccumulation metrics

When users click on the ‘Calculate and Display’ button, some results are provided by default. First, the *BCF_k_* is given as a probability distribution (**Fig. 2-a**) and summarized with its median and its 95% uncertainty limits, that is 95% credible interval delimited by the 2.5^th^ and the 97.5^th^ centiles of the posterior probability distribution (**Table 3-a**). If users ask for the *BCF_SS_*, its probability distribution is also delivered (**Fig. 2-b**) also summarized with the median and the 95% uncertainty limits (**Table 3-a**). Credible intervals are crucial information to quantify the uncertainty on parameter estimates. If data are available for several exposure routes (according to the experimental design) and uploaded within MOSAIC_bioacc_, the BCF/BSAF/BMF metrics are displayed in separate tabs (see **Annex 4** for an example).

**Figure 2.**
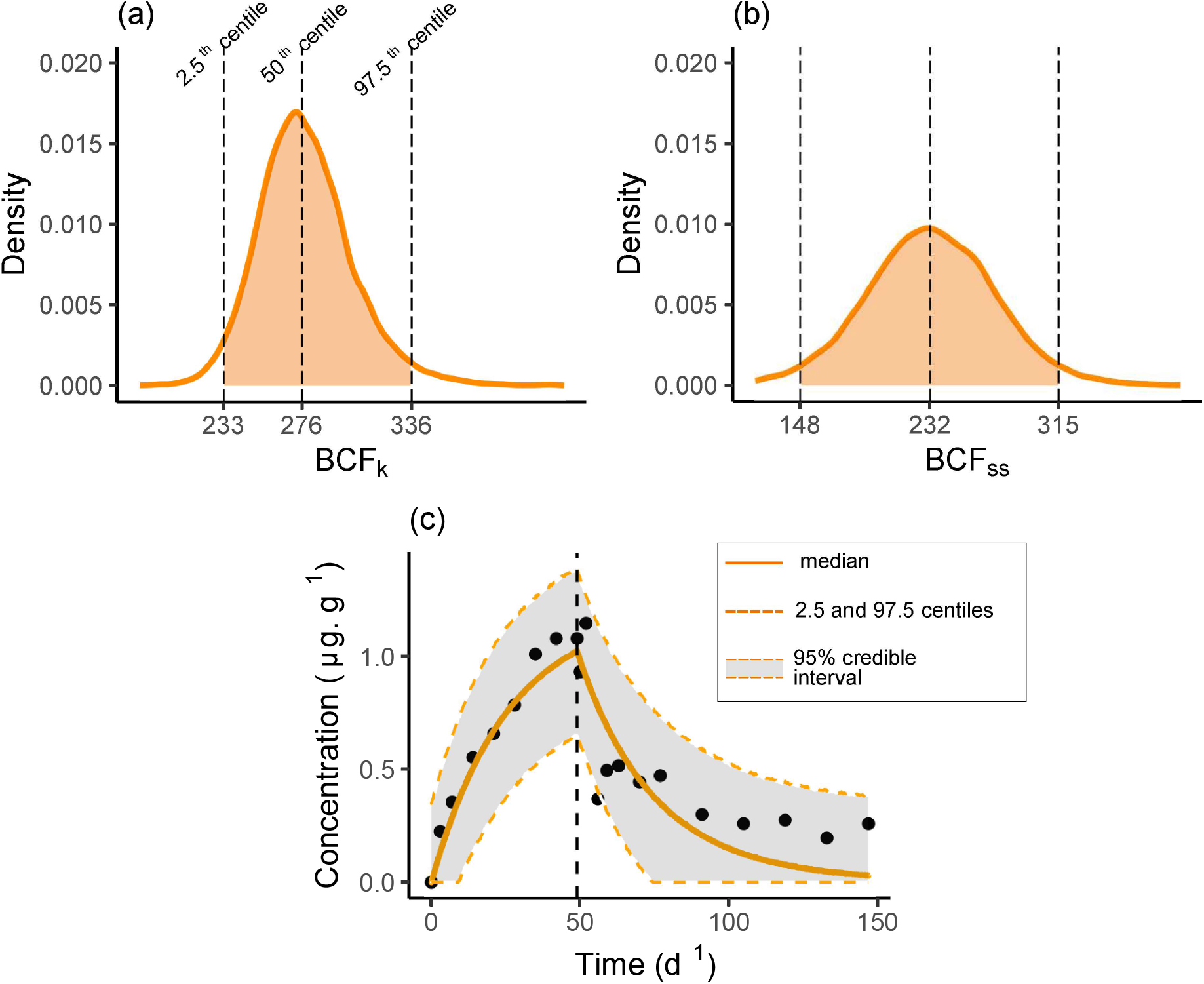
Probability distributions of *BCF_k_* (a) and *BCF_SS_* (b). The middle-dotted line represents the median value, while left and right dotted lines delimit the 2.5^th^ and 97.5^th^ centiles. (c): Observed (black dots) and predicted contaminant concentrations in the organisms (μg.g^-1^), where the median curve is displayed as the solid orange line and the uncertainty band as the grey zone, delimited by the 2.5^th^ and 97.5^th^ centiles in dotted orange lines.

**Table 3.** Example of BCF (a) and parameter (b) estimates expressed as medians (50^th^ centile) and 95% credibility intervals (2.5^th^ - 97.5^th^ centiles). Hyphens stand for dimensionless parameters.

|  | 2.5<sup>th</sup> | 50<sup>th</sup> | 97.5<sup>th</sup> | Units |
| --- | --- | --- | --- | --- |
| (a) |  |  |  |  |
| $BCF_k$ | 233 | 276 | 338 | - |
| $BCF_{ss}$ | 147 | 233 | 317 | - |
| (b) |  |  |  |  |
| $k_{uw}$ | 7.388 | 10.59 | 15.45 | d<sup>-1</sup> |
| $k_{ee}$ | 0.02307 | 0.03851 | 0.06091 | d<sup>-1</sup> |
| $\sigma_{Cp}$ | 0.1249 | 0.1684 | 0.2445 | µg.g<sup>-1</sup> |

### Predictions

Following bioaccumulation metrics calculations, the fitted curve and its uncertainty band superimposed to the observations is provided (**Fig. 2-c**). From the joint posterior distribution of model parameters, MOSAIC_bioacc_ then provides the marginal posterior distributions for each parameter, which are also summarized with quantiles in a table (**Table 3-b**): medians (for point estimates) and 2.5^th^ and 97.5^th^ centiles (for 95% credible intervals).

### Goodness-of-fit criteria

After fitting plots, several goodness-of-fit criteria follow in a prioritized order chosen based on their relevance and their ease of interpretation. The fitting quality of the model can be first checked using the Posterior Predictive Check (PPC) plot: the idea is to compare each observed value to its prediction from the fitted model at the corresponding exposure concentration associated with its 95% credible interval. If the fit is correct, we expect to get 95% of the observed values falling within the 95% credible intervals of their predictions. As shown on **Fig. 3-a**, *x*-axis locates the observed values, while the *y*-axis reports their median predictions (black dots) with their 95% credible intervals (vertical segments).

**Figure 3.**
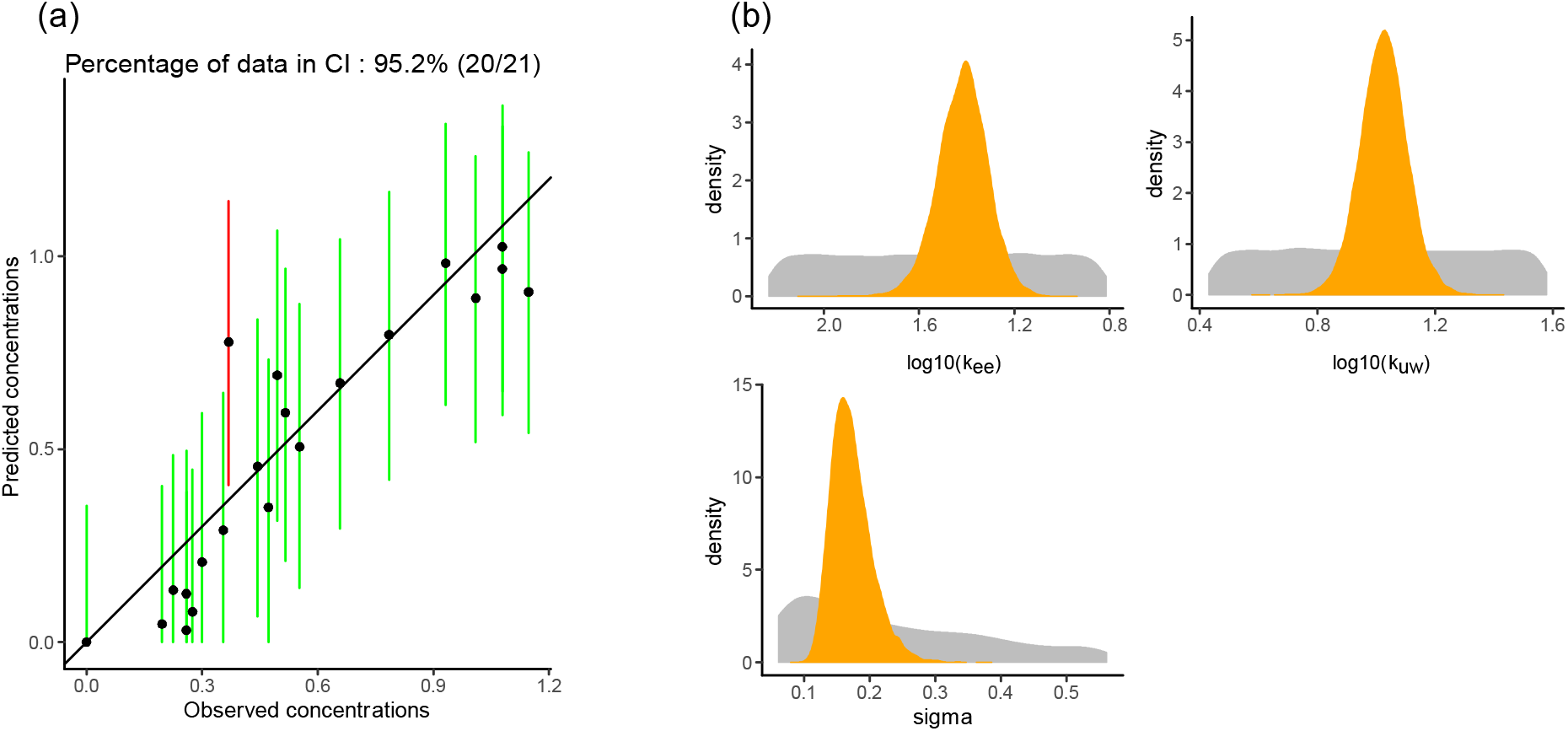
Some goodness-of-fit criteria as provided by MOSAIC_bioacc_: (a) Posterior Predictive Check (PPC) where observed values are read on the *x*-axis, while the *y*-axis reports median predictions (black dots) and their 95% credible intervals (vertical segments, coloured in green if they contain the observed value, in red otherwise); (b) prior (grey) and posterior (orange) marginal distributions of parameters of the chosen TK model (here the simplest one).

The relevance of the inference process can also be checked using the comparison of prior and posterior distributions for each model parameter. The overall expectation is to get a narrower posterior distribution compared to the prior one for each parameter, reflecting that data contributed enough to precisely estimate parameters (**Fig.3-b**). Users have the possibility to select plots for deterministic (*e.g*., *k_uw_*, *k_ee_*) or stochastic (*e.g*., *σ_C_p__*) parameters.

Then, MOSAIC_bioacc_ provides a coloured matrix in order to see at a glance the most correlated or anti-correlated parameters, in order to quickly diagnose potential problems of precision due to highly correlated parameters. Moreover, MOSAIC_bioacc_ provides plots to visualize correlations between parameters (**Fig. S1-a**). Such a plot is obtained by projecting the joint posterior distribution as a matrix in planes of parameter pairs where contours have shapes reflecting both the sign and the strength of the correlations (sub-diagonal). The correlation plot also gives marginal posterior distribution of each model parameter (diagonal) and Pearson correlation coefficients (upper diagonal). Correlations between parameters are important to consider in particular when they are high (namely, greater than 0.75) what would mean that one parameter estimate could considerably influence the other, and reciprocally. Users can display the correlation plot for deterministic parameters only or for all parameters.

The convergence of MCMC chains can be checked with the Gelman-Rubin diagnostic (Gelman and Rubin 1992) expressed via the potential scale reduction factor (PSRF) which is expected to be close to 1.00 (**Fig. S1-b**). It can be also visually verified from the MCMC trace plots, which show the time series of the sampling process leading to the posterior distribution for each parameter; it is expected to get overlap of all MCMC chains (**Fig. S1-c**). Users can visualize the MCMC trace plots for deterministic or stochastic parameters.

Finally, the Deviance Information Criterion (DIC) is provided. It is a penalized deviance statistics accounting for the number of parameters that is only useful to compare several models fitted to a same data set. Models with lower DIC values will be preferred. So, the DIC is only useful when several sub-models are compared based on different choices of parameters from all the possible combinations that the users can choose from the beginning according to the uploaded data.

### Downloads

At the bottom of the result web page, all outputs, either separately or as a full report, can be downloaded. Users can also download the entire R code corresponding to all calculations and graphs from the uploaded data set. This ensures transparency and reproducibility of all MOSAIC_bioacc_ results. This R code can be used as a steppingstone to change default options or to perform further analyses directly in the R software. For example, users can modify figures at their convenience or make several analyses on several data sets at the same time.

## CONCLUSION

Offering MOSAIC_bioacc_ as a new on-line service, free, user-friendly and ready-to-use, raises important methodological issues: (1) automation of the inference process, in particular for Bayesian inference; (2) options and choices in a transparent and facilitated way for users; (3) default outputs whose order has been chosen to facilitate their step-by-step interpretation; (4) easy accessibility to figures, tables and R code to be downloaded under different convenient formats; and (5) a final full report of all MOSAIC_bioacc_ analyses. Besides its user-friendliness, MOSAIC_bioacc_ is free of use while ensuring privacy of the uploaded data as well as transparency and reproducibility of results, together with a short response time. MOSAIC_bioacc_ is particularly useful to estimate parameters of TK models leading to predictions of chemical concentrations bioaccumulated within living organism (whatever the species, aquatic, aerial or terrestrial) from accumulation-depuration data, even standard ones. MOSAIC_bioacc_ could thus be of particular interest for risk assessors and decision makers in their daily work of evaluating dossiers, *e.g*., for market authorisation of active substances (EFSA PPR Panel 2018). Indeed, all results provided by MOSAIC_bioacc_ account for uncertainty and correlations between parameters, making possible to reproduce any previous analysis that would need to be confirmed. MOSAIC_bioacc_ can also be used for more exploratory research purposes by any environmental scientists or ecotoxicologists when accumulation-depuration data are collected and need to be analysed. MOSAIC_bioacc_ allows analyses for any species-compound combinations under consideration even with biotransformation processes, allowing users to easily perform TK analyses accounting for several exposure routes and phase I metabolites. MOSAIC_bioacc_ is available as a new statistical service dedicated to TK modelling approaches embedded within the MOSAIC platform (Charles et al. 2018). This makes MOSAIC an all-in-one facility for many applications. MOSAIC_bioacc_ will be shortly extended with a prediction tool, in order to help in designing new TK experiments, for example, when a new species-compound combination requires attention or when additional data are needed to get a better precision on bioaccumulation metrics. To gain again in generality, an R-package (rbioacc) is already in progress to include all the functionalities encompassed in MOSAIC_bioacc_, which will also make it possible to extend the use of TK models when the exposure concentrations vary over time, and when it is necessary to consider non-first order kinetics.

## Acknowledgement

The authors are thankful to ANSES for providing the financial support. The MOSAIC_bioacc_ web tool is hosted at the Rhône-Alpes Bioinformatics Center PRABI (PRABI, 2020). This work benefited from the French GDR “Aquatic Ecotoxicology” framework which aims at fostering stimulating scientific discussions and collaborations for more integrative approaches. This work is part of the ANR project APPROve (ANR-18-CE34-0013) for an integrated approach to propose proteomics for biomonitoring: accumulation, fate and multimarkers (https://anr.fr/Projet-ANR-18-CE34-0013). This work was also made under the umbrella of the Graduate School H2O’Lyon (ANR-17-EURE-0018) and “Université de Lyon” (UdL), as part of the program “Investissements d’Avenir” run by “Agence Nationale de la Recherche” (ANR). The authors are truly grateful to the anonymous colleagues who participated in testing MOSAIC_bioacc_ and for giving their feedback. The authors also are grateful for Benoît BRET for creating the logo for MOSAIC_bioacc_. The authors declare no competing interests.

## Data availability

Data are accessible directly within MOSAIC_bioacc_ at https://mosaic.univ-lyon1.fr/bioacc.

## SUPPLEMENTAL DATA

Supplemental data are available on-line.

## Notes

https://mosaic.univ-lyon1.fr/bioacc

### Competing Interest Statement

The authors have declared no competing interest.

