## supporting information v5 for "New perspectives on the calculation of bioaccumulation metrics for active substances in living organisms"

Revised Manuscript [2021-04-19]

**Contents**

**Figure S1** 2

**Annex 1:** Generic solution of TK models 3

**Annex 2:** Nested models deriving from the complete one 8

**Annex 3:** Example of a report provided by $MOSAIC_{bioacc}$ 11

**Annex 4:** Example of a report provided by $MOSAIC_{bioacc}$ with sediment exposure 26

**Annex 5:** Example of a report provided by $MOSAIC_{bioacc}$ with food exposure 38

**Annex 6:** Steady-state calculation for the simplest TK model 50

**Figure S1.** Other goodness-of-fit criteria provided by MOSAIC$_{bioacc}$: correlation plot, (b) Potential Scale Reduction Factor (PSRF) and (c) MCMC traces.

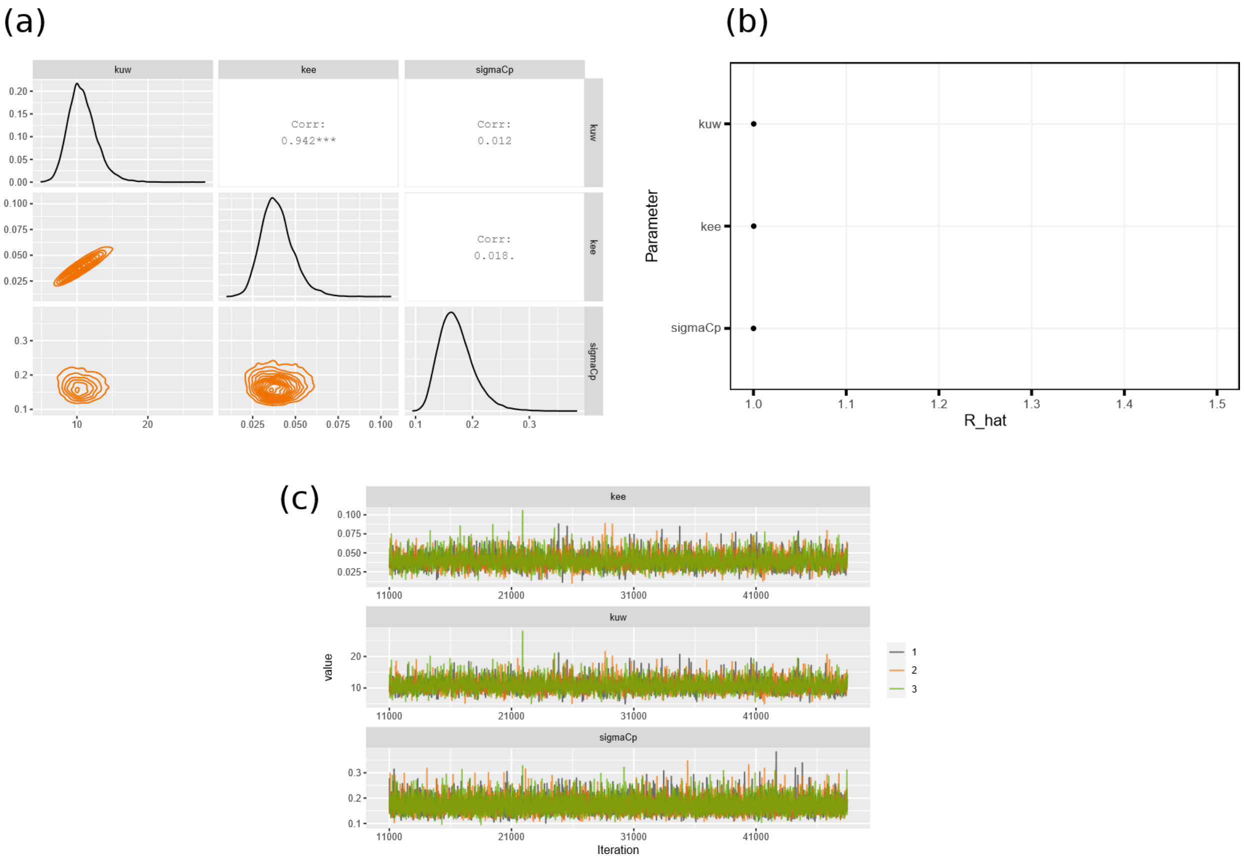

#### ANNEX 1: GENERIC SOLUTION OF TK MODELS

##### Introduction

In this document, we consider a very generic one-compartment model with several exposure sources (*e.g.*, by water, sediment and/or food), several elimination processes (*e.g.*, direct elimination, dilution by growth and/or biotransformation) and several metabolites of the parent chemical compound.

In theory, the exposure concentration may be variable over time, but in this case, there is no analytical solution of the TK model; only a numerical solution can be calculated with an appropriate algorithm. This document then assumes a constant exposure concentration for all exposure sources and provides the corresponding exact solution of the one-compartment TK model.

##### Symbols

| Symbol | Meaning |
|---|---|
| $I$ | total number of exposure sources |
| $J$ | total number of elimination processes |
| $L$ | total number of metabolites |
| $i$ | index of exposure sources, $i = 1 \dots I$ |
| $j$ | index of elimination processes, $j = 1 \dots J$ |
| $\ell$ | index of metabolites, $\ell = 1 \dots L$ |
| $t$ | time (expressed in time units) |
| $c_i$ | exposure concentration of route $i$ (in $\mu g.mL^{-1}$) |
| $C_p(t)$ | internal concentration of the parent compound at time $t$ (in $\mu g.g^{-1}$) |
| $C_{m_\ell}(t)$ | internal concentration of metabolite $\ell$ (in $\mu g.g^{-1}$) |
| $k_{u_i}$ | uptake rate of exposure source $i$ (expressed per *time*$^{-1}$) |
| $k_{e_j}$ | elimination rates of elimination process $j$ (expressed per *time*$^{-1}$) |
| $k_{e_\ell}$ | elimination rates of metabolite $\ell$ (expressed per *time*$^{-1}$) |
| $k_{m_\ell}$ | metabolization rate of metabolite $\ell$ (expressed per *time*$^{-1}$) |
| $t_c$ | duration of the accumulation phase (expressed per time units) |

##### General notations

| Letter | Meaning |
|---|---|
| $U = \sum_{i=1}^{I} k_{u_i} c_i$ | sum of all uptake terms |
| $E = \sum_{j=1}^{J} k_{e_j}$ | sum of all elimination terms for the parent compound |
| $M = \sum_{\ell=1}^{L} k_{m_\ell}$ | sum of all elimination terms for metabolite $\ell$ |

#### Intermediate notations

| Letter |
| --- |
| $R = \dfrac{U}{E + M}$ |
| $D = k_{e_\ell} - (E + M)$ |
| $Q = C_0 - R\left(1 - e^{(E+M)t_c}\right)$ |

#### Accumulation phase ($0 \leq t \leq t_c$)

##### System of ordinary differential equations

For the accumulation phase, the system of ordinary differential equations (ODE) can be written in two generic accumulation equations (AE) as follows:

$$\begin{cases} \dfrac{dC_p(t)}{dt} = U - (E + M)C_p(t) \quad (AE_1) \\ \dfrac{dC_{m_\ell}(t)}{dt} = k_{m_\ell}C_p(t) - k_{e_\ell}C_{m_\ell}(t) \quad \forall \ell = 1 \dots M \quad (AE_2) \end{cases}$$

##### Solution for the parent compound (eq. $AE_1$)

Equation ($AE_1$) is a linear first ODE with constant coefficient and a second member admitting $C_{part}(t) = \frac{U}{E+M} = R$ as a particular solution.

The solution of ($AE_1$) without the second member writes:

$$C_p(t) = Ke^{-(E+M)t} \text{ with } K \in \mathbb{R}^+$$

leading to the following general solution for equation ($AE_1$):

$$C_p(t) = Ke^{-(E+M)t} + R \text{ with } K \in \mathbb{R}^+$$

From the initial condition $C(t = 0) = C_0$ ($C_0 \geq 0$), we finally get the expression of the internal concentration of the parent compound for the accumulation phase:

$$C_p(t) = (C_0 - R)e^{-(E+M)t} + R \qquad (AS_1)$$

##### Solution for metabolite $\ell$ (eq. $AE_2$)

Equation ($AE_2$) is also a linear first ODE with constant coefficients and a second member.

The solution of ($AE_2$) without the second member writes:

$$C_{m_\ell}(t) = Ke^{-k_{e_\ell}t} \text{ with } K \in \mathbb{R}^+$$

The method know as variation of constants consists of writing the general solution of ($AE_2$) as:

$$C_{m_\ell}(t) = K(t)e^{-k_{e_\ell}t}$$

and to find function $K(t)$, by deriving and re-injecting the result into ($AE_2$). The derivative writes:

$$\frac{dC_{m_\ell}(t)}{dt} = \frac{dK(t)}{dt}e^{-k_{e_\ell}t} - K(t)k_{e_\ell}e^{-k_{e_\ell}t}$$

The re-injection into $(AE_2)$ leads to:

$$\frac{dK(t)}{dt} = k_{m_\ell}\left((C_0 - R)e^{Dt} + Re^{k_{e_\ell}t}\right)$$

which integrates into:

$$K(t) = k_{m_\ell}\left(\frac{C_0 - R}{D}e^{Dt} + \frac{R}{k_{e_\ell}}e^{k_{e_\ell}t}\right) + C \text{ with } C \in \mathbb{R}$$

The general solution of $(AE_2)$ finally writes as follows:

$$C_{m_\ell}(t) = k_{m_\ell}\left(\frac{C_0 - R}{D}e^{-(E+M)t} + \frac{R}{k_{e_\ell}}\right) + Ce^{-k_{e_\ell}t} \text{ with } C \in \mathbb{R}$$

From the initial condition $C_{m_\ell}(t = 0) = 0$, we finally get the expression of the internal concentration of metabolite $\ell$ for the accumulation phase:

$$C_{m_\ell}(t) = k_{m_\ell}\left(\frac{C_0 - R}{D}\left(e^{-(E+M)t} - e^{-k_{e_\ell}t}\right) + \frac{R}{k_{e_\ell}}\left(1 - e^{-k_{e_\ell}t}\right)\right) \qquad (AS_2)$$

#### Depuration phase ($t > t_c$)

##### System of ordinary differential equations

For the depuration phase, the system of ODE can be written in two generic depuration equations (DE) as follows:

$$\begin{cases} \dfrac{dC_p(t)}{dt} = -(E + M)C_p(t) & (DE_1) \\ \dfrac{dC_{m_\ell}(t)}{dt} = k_{m_\ell}C_p(t) - k_{e_\ell}C_{m_\ell}(t) \quad \forall \ell = 1 \dots M & (DE_2) \end{cases}$$

##### Solution for the parent compound (eq. $DE_1$)

Equation $(DE_1)$ has a general solution of the following form:

$$C_p(t) = Ke^{-(E+M)t}$$

For the depuration phase and the parent compound, the initial condition comes the calculation of the internal parent compound concentration at the end of the accumulation (*i.e.*, $t = t_c$) from solution $(AS_1)$, that is:

$$C_p(t_c) = (C_0 - R)e^{-(E+M)t_c} + R$$

From the general solution of $(DE_1)$, we get $C_p(t_c) = Ke^{-(E+M)t_c}$ leading to:

$$K = C_0 - R + Re^{(E+M)t_c}$$

Then, the final expression of the internal concentration of the parent compound for the depuration phase writes:

$$C_p(t) = \left(C_0 - R\left(1 - e^{(E+M)t_c}\right)\right)e^{-(E+M)t} \qquad (DS_1)$$

**Solution for metabolite $\ell$ (Eq. $DE_2$)**

Temporarily, solution $(DS_1)$ above can be written as $C_p(t) = Qe^{-(E+M)t}$ with constant $Q$ defined at the beginning of the document.

Equation $(DE_2)$ is a linear ODE of first order with constant coefficients and a second member.

The solution of the equation without the second member writes:

$$C_{m_\ell}(t) = Ke^{-k_{e_\ell}t}$$

As earlier, we use the variation of constants method by writing the general solution of $(DE_2)$ as $C_{m_\ell}(t) = K(t)e^{-k_{e_\ell}t}$ and searching for function $K(t)$.

The derivative of $C_{m_\ell}(t)$ is $\frac{dK(t)}{dt}e^{-k_{e_\ell}t} - k_{e_\ell}K(t)e^{-k_{e_\ell}t}$.

The re-injection of this derivative into $(DE_2)$ leads to:

$$\frac{dK(t)}{dt} = k_{m_\ell}Qe^{Dt}$$

which integrates into:

$$K(t) = k_{m_\ell}\frac{Q}{D}e^{Dt} + C \text{ with } C \in \mathbb{R}$$

finally leading to the general solution of $(DE_2)$:

$$C_{m_\ell}(t) = k_{m_\ell}\frac{Q}{D}e^{-(E+M)t} + Ce^{-k_{e_\ell}t}$$

Constant $C$ is determined with the initial condition, i.e., the internal concentration of metabolite $\ell$ at $t = t_c$ both at the end of the accumulation phase and at the beginning of the depuration phase.

From the previous equation, we get $C_{m_\ell}(t_c) = k_{m_\ell}\frac{Q}{D}e^{-(E+M)t_c} + Ce^{-k_{e_\ell}t_c}$.

From solution $(AS_2)$, we get:

$$C_{m_\ell}(t_c) = k_{m_\ell}\left(\frac{C_0 - R}{D}\left(e^{-(E+M)t_c} - e^{-k_{e_\ell}t_c}\right) + \frac{R}{k_{e_\ell}}\left(1 - e^{-k_{e_\ell}t_c}\right)\right)$$

We finally get the following expression for constant $C$:

$$C = k_{m_\ell}\left(\frac{R}{k_{e_\ell}}\left(e^{k_{e_\ell}t_c} - 1\right) - \frac{C_0 - R}{D} - \frac{Q}{D}e^{k_{e_\ell}t_c}\right)$$

Replacing constant C gives the final expression of the internal concentration of metabolite $\ell$ for the depuration phase:

$$C_{m_\ell}(t) = k_{m_\ell}\left(\frac{Q}{D}e^{-(E+M)t} + \frac{R}{k_{e_\ell}}\left(e^{-k_{e_\ell}(t-t_c)} - e^{-k_{e_\ell}t}\right) - \frac{C_0 - R}{D}e^{-k_{e_\ell}t} - \frac{Q}{D}e^{-k_{e_\ell}(t-t_c)}\right)$$

Replacing constant $Q$ by its own expression gives:

$$C_{m_\ell}(t) = k_{m_\ell}\left(\frac{C_0 - R}{D}\left(e^{-(E+M)t} - e^{-k_{e_\ell}t}\right) + \frac{R}{k_{e_\ell}}\left(e^{-k_{e_\ell}(t-t_c)} - e^{-k_{e_\ell}t}\right) + \frac{R}{D}\left(e^{-k_{e_\ell}(t-t_c)} - e^{-k_{e_\ell}(t-t_c)}\right)\right) \qquad (DS_2)$$

**Final set of solutions for both phases**

Reminding the following intermediate notations $R = \frac{U}{E+M}$ and $D = k_{e_\ell} - (E+M)$ we can obtain the final set of solutions for both phases:

- Internal concentration of the parent compound for the accumulation phase:

$$C_p(t) = \left(C_0 - \frac{U}{E+M}\right)e^{-(E+M)t} + \frac{U}{E+M} \qquad (AS_1) \quad \forall 0 \leq t \leq t_c$$

- Internal concentration of metabolite $\ell$ for the acummulation phase:

$$C_{m_\ell}(t) = k_{m_\ell}\left(\frac{C_0(E+M) - U}{(E+M)\big(k_{e_\ell} - (E+M)\big)}\left(e^{-(E+M)t} - e^{-k_{e_\ell}t}\right) + \frac{U}{k_{e_\ell}(E+M)}\left(1 - e^{-k_{e_\ell}t}\right)\right) \qquad (AS_2)$$

- Internal concentration of the parent compound for the depuration phase writes:

$$C_p(t) = \left(C_0 - \frac{U}{E+M}\left(1 - e^{(E+M)t_c}\right)\right)e^{-(E+M)t} \qquad (DS_1) \quad \forall t > t_c$$

- Internal concentration of metabolite $\ell$ for the depuration phase:

$$C_{m_\ell}(t) = \frac{k_{m_\ell}C_0(E+M) - U}{(E+M)(k_{e_\ell} - (E+M))}\left(e^{-(E+M)t} - e^{-k_{e_\ell}t}\right) + \frac{k_{m_\ell}U}{k_{e_\ell}(E+M)}\left(e^{-k_{e_\ell}(t-t_c)} - e^{-k_{e_\ell}t}\right) + \frac{k_{m_\ell}U}{(E+M)(k_{e_\ell} - (E+M))}\left(e^{-k_{e_\ell}(t-t_c)} - e^{-k_{e_\ell}(t-t_c)}\right) \qquad (DS_2) \quad \forall t > t_c$$

In the end, we could replace constants $U$, $E$ and $M$ by $U = \sum_{i=1}^{I} k_{u_i} c_i$, $E = \sum_{j=1}^{J} k_{e_j}$ and $M = \sum_{\ell=1}^{L} k_{m_\ell}$, respectively, in order to write the very final set of solutions with all the parameters to estimate from observed data with an inference process.

#### ANNEX 2: NESTED MODELS DERIVING FROM THE COMPLETE ONE

Within the $MOSAIC_{bioacc}$ web app, we choose to only propose four routes of exposure ($I$ = 4, *i.e.*, water, pore water, food and sediment), three processes of elimination ($J$ = 3, *i.e.*, excretion, growth dilution and biotransformation) where all metabolites directly come from the parent compound. When three metabolites are considered ($L$ = 3), this can be expressed as follows (Eqs. ($SE1$) to ($SE5$)):

$$\begin{cases} \frac{dC_p(t)}{dt} = (k_{u_w} c_w) + \left(k_{u_{pw}} c_{pw}\right) + (k_{u_s} c_s) + \left(k_{u_f} c_f\right) - \left(k_{e_e} + k_{e_g} + k_{m_1} + k_{m_2} + k_{m_3}\right) C_p(t) & (SE_1) \\ \frac{dC_{m_1}(t)}{dt} = k_{m_1} C_p(t) - k_{e_{m1}} C_{m_1}(t) & (SE_2) \\ \frac{dC_{m_2}(t)}{dt} = k_{m_2} C_p(t) - k_{e_{m2}} C_{m_2}(t) & (SE_3) \\ \frac{dC_{m_3}(t)}{dt} = k_{m_3} C_p(t) - k_{e_{m3}} C_{m_3}(t) & (SE_4) \end{cases}$$

$$\frac{dG(t)}{dt} = k_{e_g} \times (g_{max} - G(t)) \qquad (SE_5)$$

Parameters are summarized in the following table:

| Symbol | Meaning |
|---|---|
| $t$ | time (expressed in time unit) |
| $c_w$ | water exposure concentration (in $\mu g.mL^{-1}$) |
| $c_{pw}$ | pore water exposure concentration (in $\mu g.mL^{-1}$) |
| $c_f$ | food exposure concentration (in $\mu g.g^{-1}$) |
| $c_s$ | sediment exposure concentration (in $\mu g.g^{-1}$) |
| $C_p(t)$ | internal concentration of the parent compound at time $t$ (in $\mu g.g^{-1}$) |
| $C_{m1}(t)$ | internal concentration of metabolite 1 (in $\mu g.g^{-1}$) |
| $C_{m2}(t)$ | internal concentration of metabolite 2 (in $\mu g.g^{-1}$) |
| $C_{m3}(t)$ | internal concentration of metabolite 3 (in $\mu g.g^{-1}$) |
| $k_{uw}$ | uptake rate from water (expressed in $time^{-1}$) |
| $k_{upw}$ | uptake rate from pore water (expressed in $time^{-1}$) |
| $k_{uf}$ | uptake rate from food (expressed in $time^{-1}$) |
| $k_{us}$ | uptake rate from sediment (expressed in $time^{-1}$) |
| $k_{ee}$ | elimination rate by excretion (expressed in $time^{-1}$) |
| $k_{eg}$ | elimination rate by growth dilution (expressed in $time^{-1}$) |
| $k_{em1}$ | elimination rate of metabolite 1 (expressed in $time^{-1}$) |
| $k_{em2}$ | elimination rate of metabolite 2 (expressed in $time^{-1}$) |
| $k_{em3}$ | elimination rate of metabolite 3 (expressed in $time^{-1}$) |
| $k_{m1}$ | metabolization rate of metabolite 1 (expressed in $time^{-1}$) |
| $k_{m2}$ | metabolization rate of metabolite 2 (expressed in $time^{-1}$) |
| $k_{m3}$ | metabolization rate of metabolite 3 (expressed in $time^{-1}$) |
| $t_c$ | duration of the accumulation phase (expressed per time unit) |
| $G(t)$ | growth of the organism at time $t$ (expressed per $g$) |
| $g_{max}$ | asymptotic organism growth (expressed per $g$) |

We assume a Gaussian probability distribution for the contaminant and its metabolites concentration within the organism, as well as for the growth of the organism, as follow (Eqs. ($SE_{10}$) to ($SE_{14}$)):

$$C_{obs,p}(t) \sim \mathcal{N}\left(C_p(t), \sigma_{C_p}^2\right) \quad (SE_{10})$$

$$C_{obs,m1}(t) \sim \mathcal{N}\left(C_{m_1}(t), \sigma_{m1}^2\right) \quad (SE_{11})$$

$$C_{obs,m2}(t) \sim \mathcal{N}\left(C_{m_2}(t), \sigma_{m2}^2\right) \quad (SE_{12})$$

$$C_{obs,m3}(t) \sim \mathcal{N}\left(C_{m_3}(t), \sigma_{m3}^2\right) \quad (SE_{13})$$

$$G_{obs}(t) \sim \mathcal{N}(G(t), \sigma_G^2) \quad (SE_{14})$$

Where:

| Symbol | Meaning |
|---|---|
| $\mathcal{N}$ | Normal probability distribution |
| $C_{obs,p}(t)$ | internal measured concentration of the parent compound at time $t$ (in $\mu g.g^{-1}$) |
| $C_{obs,m_1}(t)$ | internal measured concentration of metabolite 1 (in $\mu g.g^{-1}$) |
| $C_{obs,m_2}(t)$ | internal measured concentration of metabolite 2 (in $\mu g.g^{-1}$) |
| $C_{obs,m_3}(t)$ | internal measured concentration of metabolite 3 (in $\mu g.g^{-1}$) |
| $G_{obs}(t)$ | measured growth of the organism at time $t$ (in $\mu g.g^{-1}$) |
| $\sigma_{C_p}$ | standard deviation for the internal concentration of the parent compound (in $\mu g.g^{-1}$) |
| $\sigma_{m_1}$ | standard deviation for the internal concentration of metabolite 1 (in $\mu g.g^{-1}$) |
| $\sigma_{m_2}$ | standard deviation for the internal concentration of metabolite 2 (in $\mu g.g^{-1}$) |
| $\sigma_{m_3}$ | standard deviation for the internal concentration of metabolite 3 (in $\mu g.g^{-1}$) |
| $\sigma_G$ | standard deviation for the growth of the organism (in $g$) |

Hence, the maximum number of parameters is 19 when the model considers three metabolites, and the user will have at the end a total of 112 possible models depending on the combination it chooses (Table S1).

**Table S1**. Example of parameters combinations available in $MOSAIC_{bioacc}$ web app to perform calculations according to the biological processes the user considers in its data. A value of 1 stands for the parameter, 0 for not including it, according to the chosen processes.

| kuw | kus | kuf | kupw | km1 | km2 | km3 | kee | keg | kem1 | kem2 | kem3 | *g*0 | gmax | $\sigma C_p$ | $\sigma_G$ | σm1 | σm2 | σm3 |
|---|---|---|---|---|---|---|---|---|---|---|---|---|---|---|---|---|---|---|
| 1 | 0 | 0 | 0 | 0 | 0 | 0 | 1 | 0 | 0 | 0 | 0 | 0 | 0 | 1 | 0 | 0 | 0 | 0 |
| 1 | 0 | 0 | 0 | 0 | 0 | 0 | 1 | 1 | 0 | 0 | 0 | 1 | 1 | 1 | 1 | 0 | 0 | 0 |
| 1 | 0 | 0 | 0 | 1 | 0 | 0 | 1 | 1 | 1 | 0 | 0 | 1 | 1 | 1 | 1 | 1 | 0 | 0 |
| 1 | 0 | 0 | 0 | 1 | 1 | 0 | 1 | 1 | 1 | 1 | 0 | 1 | 1 | 1 | 1 | 1 | 1 | 0 |
| 1 | 0 | 0 | 0 | 1 | 1 | 1 | 1 | 1 | 1 | 1 | 1 | 1 | 1 | 1 | 1 | 1 | 1 | 1 |
| 1 | 1 | 0 | 0 | 0 | 0 | 0 | 1 | 0 | 0 | 0 | 0 | 0 | 0 | 1 | 0 | 0 | 0 | 0 |
| 1 | 1 | 0 | 0 | 0 | 0 | 0 | 1 | 1 | 0 | 0 | 0 | 1 | 1 | 1 | 1 | 0 | 0 | 0 |
| 1 | 1 | 0 | 0 | 1 | 0 | 0 | 1 | 1 | 1 | 0 | 0 | 1 | 1 | 1 | 1 | 1 | 0 | 0 |
| 1 | 1 | 0 | 0 | 1 | 1 | 0 | 1 | 1 | 1 | 1 | 0 | 1 | 1 | 1 | 1 | 1 | 1 | 0 |
| 1 | 1 | 0 | 0 | 1 | 1 | 1 | 1 | 1 | 1 | 1 | 1 | 1 | 1 | 1 | 1 | 1 | 1 | 1 |
| 1 | 1 | 1 | 0 | 0 | 0 | 0 | 1 | 0 | 0 | 0 | 0 | 0 | 0 | 1 | 0 | 0 | 0 | 0 |
| 1 | 1 | 1 | 0 | 0 | 0 | 0 | 1 | 1 | 0 | 0 | 0 | 1 | 1 | 1 | 1 | 0 | 0 | 0 |
| 1 | 1 | 1 | 0 | 1 | 0 | 0 | 1 | 1 | 1 | 0 | 0 | 1 | 1 | 1 | 1 | 1 | 0 | 0 |
| 1 | 1 | 1 | 0 | 1 | 1 | 0 | 1 | 1 | 1 | 1 | 0 | 1 | 1 | 1 | 1 | 1 | 1 | 0 |
| 1 | 1 | 1 | 0 | 1 | 1 | 1 | 1 | 1 | 1 | 1 | 1 | 1 | 1 | 1 | 1 | 1 | 1 | 1 |
| 0 | 1 | 0 | 0 | 0 | 0 | 0 | 1 | 0 | 0 | 0 | 0 | 0 | 0 | 1 | 0 | 0 | 0 | 0 |
| 0 | 1 | 0 | 0 | 0 | 0 | 0 | 1 | 1 | 0 | 0 | 0 | 1 | 1 | 1 | 1 | 0 | 0 | 0 |
| 0 | 1 | 0 | 0 | 1 | 0 | 0 | 1 | 1 | 1 | 0 | 0 | 1 | 1 | 1 | 1 | 1 | 0 | 0 |
| 0 | 1 | 0 | 0 | 1 | 1 | 0 | 1 | 1 | 1 | 1 | 0 | 1 | 1 | 1 | 1 | 1 | 1 | 0 |
| 0 | 1 | 0 | 0 | 1 | 1 | 1 | 1 | 1 | 1 | 1 | 1 | 1 | 1 | 1 | 1 | 1 | 1 | 1 |
| 0 | 0 | 1 | 0 | 0 | 0 | 0 | 1 | 0 | 0 | 0 | 0 | 0 | 0 | 1 | 0 | 0 | 0 | 0 |
| 0 | 0 | 1 | 0 | 0 | 0 | 0 | 1 | 1 | 0 | 0 | 0 | 1 | 1 | 1 | 1 | 0 | 0 | 0 |
| 0 | 0 | 1 | 0 | 1 | 0 | 0 | 1 | 1 | 1 | 0 | 0 | 1 | 1 | 1 | 1 | 1 | 0 | 0 |
| 0 | 0 | 1 | 0 | 1 | 1 | 0 | 1 | 1 | 1 | 1 | 0 | 1 | 1 | 1 | 1 | 1 | 1 | 0 |
| 0 | 0 | 1 | 0 | 1 | 1 | 1 | 1 | 1 | 1 | 1 | 1 | 1 | 1 | 1 | 1 | 1 | 1 | 1 |
| 1 | 0 | 1 | 0 | 0 | 0 | 0 | 1 | 0 | 0 | 0 | 0 | 0 | 0 | 1 | 0 | 0 | 0 | 0 |
| 1 | 0 | 1 | 0 | 0 | 0 | 0 | 1 | 1 | 0 | 0 | 0 | 1 | 1 | 1 | 1 | 0 | 0 | 0 |
| 1 | 0 | 1 | 0 | 1 | 0 | 0 | 1 | 1 | 1 | 0 | 0 | 1 | 1 | 1 | 1 | 1 | 0 | 0 |
| 1 | 0 | 1 | 0 | 1 | 1 | 0 | 1 | 1 | 1 | 1 | 0 | 1 | 1 | 1 | 1 | 1 | 1 | 0 |
| 1 | 0 | 1 | 0 | 1 | 1 | 1 | 1 | 1 | 1 | 1 | 1 | 1 | 1 | 1 | 1 | 1 | 1 | 1 |
| 0 | 1 | 1 | 0 | 0 | 0 | 0 | 1 | 0 | 0 | 0 | 0 | 0 | 0 | 1 | 0 | 0 | 0 | 0 |
| 0 | 1 | 1 | 0 | 0 | 0 | 0 | 1 | 1 | 0 | 0 | 0 | 1 | 1 | 1 | 1 | 0 | 0 | 0 |
| 0 | 1 | 1 | 0 | 1 | 0 | 0 | 1 | 1 | 1 | 0 | 0 | 1 | 1 | 1 | 1 | 1 | 0 | 0 |
| 0 | 1 | 1 | 0 | 1 | 1 | 0 | 1 | 1 | 1 | 1 | 0 | 1 | 1 | 1 | 1 | 1 | 1 | 0 |
| 0 | 1 | 1 | 0 | 1 | 1 | 1 | 1 | 1 | 1 | 1 | 1 | 1 | 1 | 1 | 1 | 1 | 1 | 1 |
| 1 | 1 | 1 | 0 | 1 | 0 | 0 | 1 | 0 | 1 | 0 | 0 | 0 | 0 | 1 | 0 | 1 | 0 | 0 |
| 1 | 1 | 1 | 0 | 1 | 1 | 0 | 1 | 0 | 1 | 1 | 0 | 0 | 0 | 1 | 0 | 1 | 1 | 0 |
| 1 | 1 | 1 | 0 | 1 | 1 | 1 | 1 | 0 | 1 | 1 | 1 | 0 | 0 | 1 | 0 | 1 | 1 | 1 |
| 1 | 1 | 0 | 0 | 1 | 0 | 0 | 1 | 0 | 1 | 0 | 0 | 0 | 0 | 1 | 0 | 1 | 0 | 0 |
| 1 | 1 | 0 | 0 | 1 | 1 | 0 | 1 | 0 | 1 | 1 | 0 | 0 | 0 | 1 | 0 | 1 | 1 | 0 |
| 1 | 1 | 0 | 0 | 1 | 1 | 1 | 1 | 0 | 1 | 1 | 1 | 0 | 0 | 1 | 0 | 1 | 1 | 1 |
| 1 | 0 | 1 | 0 | 1 | 0 | 0 | 1 | 0 | 1 | 0 | 0 | 0 | 0 | 1 | 0 | 1 | 0 | 0 |
| 1 | 0 | 1 | 0 | 1 | 1 | 0 | 1 | 0 | 1 | 1 | 0 | 0 | 0 | 1 | 0 | 1 | 1 | 0 |
| 1 | 0 | 1 | 0 | 1 | 1 | 1 | 1 | 0 | 1 | 1 | 1 | 0 | 0 | 1 | 0 | 1 | 1 | 1 |
| 0 | 1 | 1 | 0 | 1 | 0 | 0 | 1 | 0 | 1 | 0 | 0 | 0 | 0 | 1 | 0 | 1 | 0 | 0 |
| 0 | 1 | 1 | 0 | 1 | 1 | 0 | 1 | 0 | 1 | 1 | 0 | 0 | 0 | 1 | 0 | 1 | 1 | 0 |
| 0 | 1 | 1 | 0 | 1 | 1 | 1 | 1 | 0 | 1 | 1 | 1 | 0 | 0 | 1 | 0 | 1 | 1 | 1 |
| 1 | 0 | 0 | 0 | 1 | 0 | 0 | 1 | 0 | 1 | 0 | 0 | 0 | 0 | 1 | 0 | 1 | 0 | 0 |
| 1 | 0 | 0 | 0 | 1 | 1 | 0 | 1 | 0 | 1 | 1 | 0 | 0 | 0 | 1 | 0 | 1 | 1 | 0 |
| 1 | 0 | 0 | 0 | 1 | 1 | 1 | 1 | 0 | 1 | 1 | 1 | 0 | 0 | 1 | 0 | 1 | 1 | 1 |
| 0 | 0 | 1 | 0 | 1 | 0 | 0 | 1 | 0 | 1 | 0 | 0 | 0 | 0 | 1 | 0 | 1 | 0 | 0 |
| 0 | 0 | 1 | 0 | 1 | 1 | 0 | 1 | 0 | 1 | 1 | 0 | 0 | 0 | 1 | 0 | 1 | 1 | 0 |
| 0 | 0 | 1 | 0 | 1 | 1 | 1 | 1 | 0 | 1 | 1 | 1 | 0 | 0 | 1 | 0 | 1 | 1 | 1 |

#### ANNEX 3: EXAMPLE OF A REPORT PROVIDED BY $\mathbf{MOSAIC_{bioacc}}$

In order to illustrate $MOSAIC_{bioacc}$ with a more complex TK model, the example file 'Male_Gammarus_seanine.csv' was selected in the application. In this example, male *Gammarus pulex* were exposed to seanine spiked water and a single exposure concentration was tested. The duration of the accumulation phase is 1.417 days. Three metabolites were quantified, and the growth of organism was included.

After calculations, the corresponding report is directly downloaded in $MOSAIC_{bioacc}$.

### MOSAIC$_{bioacc}$ REPORT
# 2020-11-03

---

This report is provided by the MOSAICbioacc application available here:

https://mosaic.univ-lyon1.fr/bioacc

MOSAIC$_{bioacc}$ uses the JAGS (version 4.3.0) and R (version 4.0.2) software, and in particular packages RJags (version 4.10) and Shiny (version 1.5.0).

The MOSAIC$_{bioacc}$ application is a turn-key web tool providing bioaccumulation metrics (BCF/BSAF/BMF) from a toxicokinetic (TK) model fitted to accumulation-depuration data. It is designed to fulfil the requirements of regulators when examining applications for market authorization of active substances.

---

###### Data summary

File used: ashauer_seanine_growth.csv

Exposure: 15.533 $\mu g.mL-^{1}$

Accumulation phase duration: 1.417 days

Number of replicates: 2

Times: 0, 0.207, 0.208, 0.52, 0.521, 0.978, 0.979, 1.207, 1.208, 1.416, 1.417, 1.999, 2, 2.978, 2.979, 3.999, 4,

4.999, 5, 5.999, 6

Exposure routes: water

Elimination routes: excretion growth biotransformation

###### Bayesian inference

Three MCMC chains were used to estimate model parameters.

Number of iterations: 273458

Thin: 73

#### TK Model

The TK model used for these calculations was:

$$\frac{dC_p(t)}{dt} = k_{uw} \times c_w - (k_{ee}+k_{eg}+k_{m1}+k_{m2}+k_{m3}) \times C_p(t) \quad \text{for } 0 \leq t \leq t_c$$

$$\frac{dC_p(t)}{dt} = -(k_{ee}+k_{eg}+k_{m1}+k_{m2}+k_{m3}) \times C_p(t) \quad \text{for } t > t_c$$

$$\frac{dC_{m1}(t)}{dt} = k_{m1} \times C_p(t) - k_{em1} \times C_{m1}(t)$$

$$\frac{dC_{m2}(t)}{dt} = k_{m2} \times C_p(t) - k_{em2} \times C_{m2}(t)$$

$$\frac{dC_{m3}(t)}{dt} = k_{m3} \times C_p(t) - k_{em3} \times C_{m3}(t)$$

$$\frac{dG(t)}{dt} = k_{eg} \times (g_{max} - G(t))$$

with:

t: time (expressed in days)

$t_c$: duration of the accumulation phase (expressed in days)

$C_p(t)$: internal concentration of the parent compound at time (expressed in $\mu g. g^{-1}$)

$k_{ee}$: elimination rates of excretion (expressed per day)

$c_w$: exposure concentration of water route (expressed in $\mu g. mL^{-1}$)

$k_{uw}$: uptake rate of water exposure (expressed per day)

$C_{m\ell}(t)$: internal concentration of metabolite $\ell$ (expressed in $\mu g. g^{-1}$)

$\ell$: index of metabolites, $\ell = 1 \ldots L$ with $L$ total number of metabolites

$k_{m\ell}$: metabolization rate of metabolite $\ell$ (expressed per day)

$k_{em\ell}$: elimination rates of metabolite $\ell$ (expressed per day)

G(t): Growth data according time $t$ (expressed in g)

$k_{eg}$: elimination rates of growth (expressed per day)

$g_{max}$: asymptotic weight at which growth is null (expressed per g)

$g_0$: weight at birth (expressed per g)

#### Bioaccumulation metrics calculation

##### Calculations

$$BCF_k = \frac{k_{uw}}{k_{ee} + k_{eg} + k_{m1} + k_{m2} + k_{m3}}$$

$$BCF_{ss} = \frac{C_p(t_c)}{c_w}$$

##### Bioconcentration factor (BCF)

##### $BCF_k$ plot

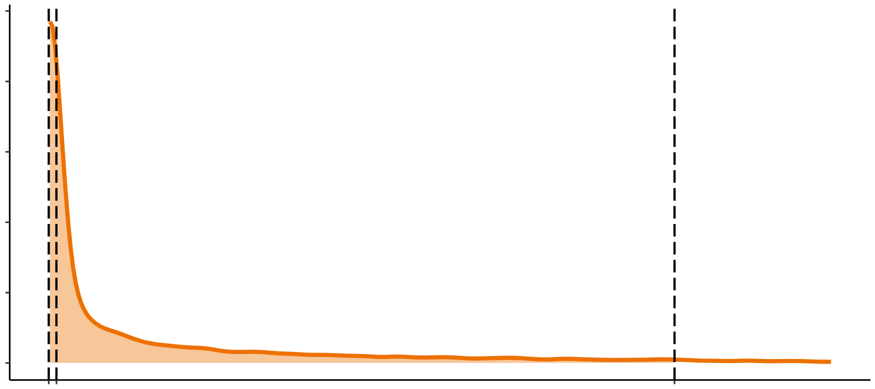

##### $BCF_{ss}$ plot

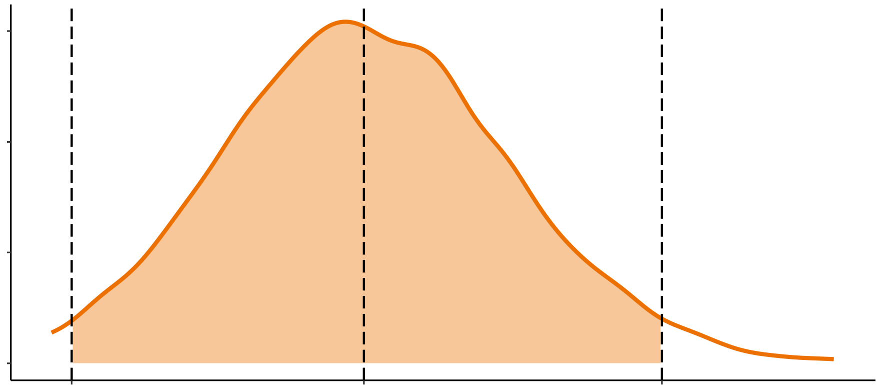

##### BCF summary

Calculated bioaccumulation factor

| | 2.5% | 50% | 97.5% | CV |
|---|---|---|---|---|
| BCFk | 62 | 3188 | 257557 | 20 |
| BCFss | 19 | 71 | 124 | 0.37 |

#### Fitting results

##### Fit plot

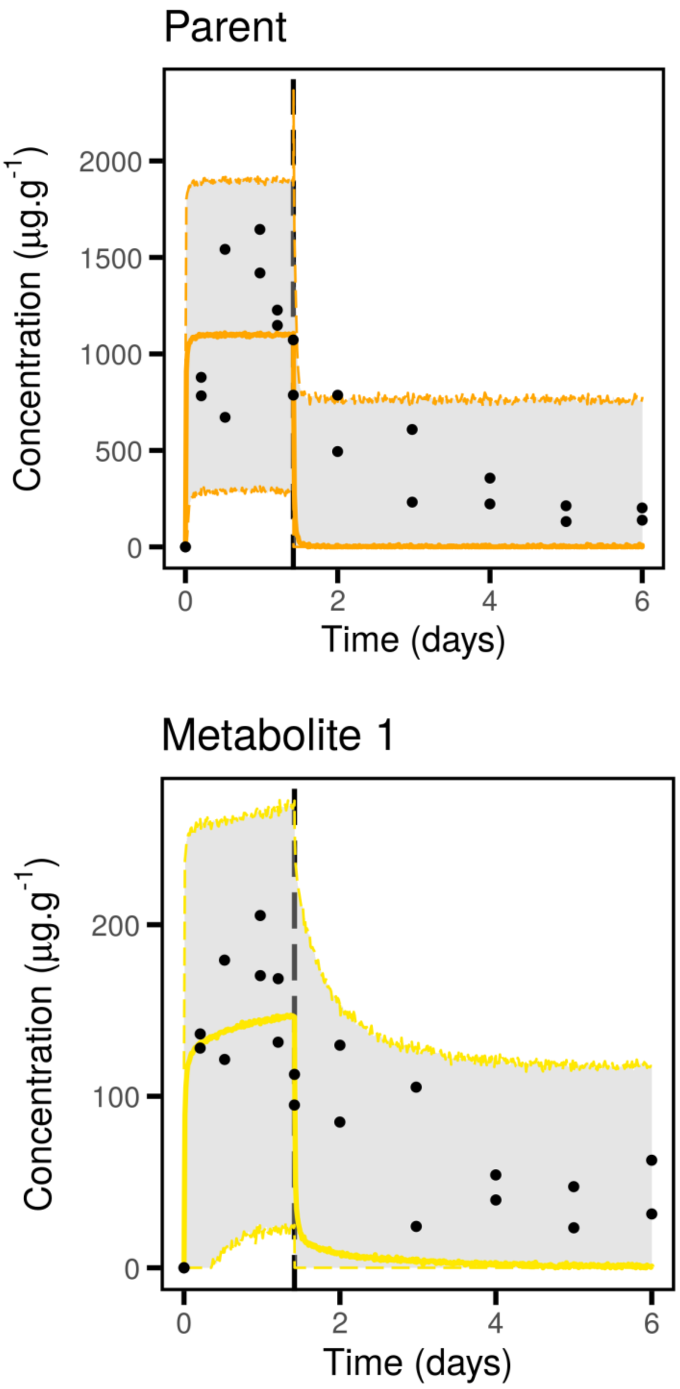

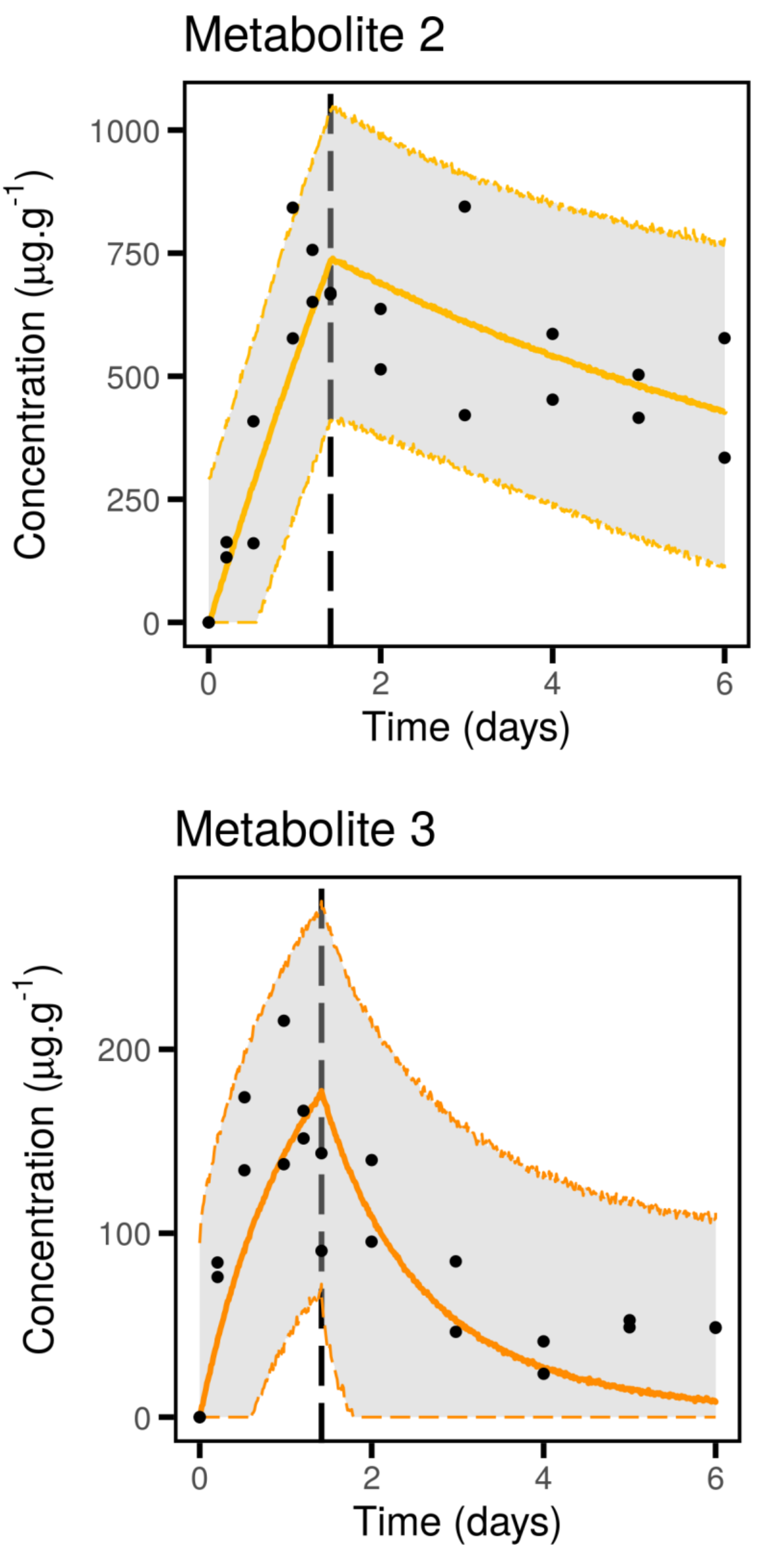
Metabolite 2
Concentration (µg.g$^{-1}$)
1000
750
500
250
0
0
2
4
6
Time (days)
Metabolite 3
Concentration (µg.g$^{-1}$)
200
100
0
0
2
4
6
Time (days)

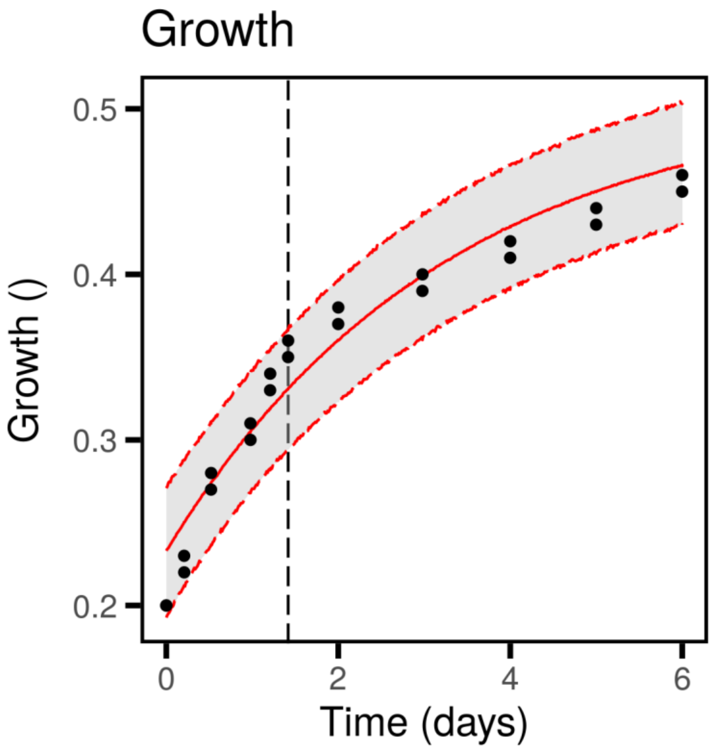


**Quantiles of estimated parameters**

| | 2.5% | 50% | 97.5% | |
|---|---|---|---|---|
| $k_{uw}$ | 1202 | 18940 | 92730 | $d^{-1}$ |
| $k_{ee}$ | 1.828e-05 | 5.086 | 1095 | $d^{-1}$ |
| $k_{eg}$ | 0.2263 | 0.3104 | 0.3735 | $d^{-1}$ |
| $k_{m1}$ | 0.1591 | 94.21 | 1246 | $d^{-1}$ |
| $k_{m2}$ | 0.3719 | 0.5176 | 0.7065 | $d^{-1}$ |
| $k_{m3}$ | 0.116 | 0.1949 | 0.8331 | $d^{-1}$ |
| $k_{em1}$ | 0.604 | 728.5 | 9604 | $d^{-1}$ |
| $k_{em2}$ | 2.447e-03 | 0.1221 | 0.2144 | $d^{-1}$ |
| $k_{em3}$ | 0.3218 | 0.7776 | 5.974 | $d^{-1}$ |
| $g_{max}$ | 0.5002 | 0.5067 | 0.5488 | $g$ |
| $g_0$ | 0.2128 | 0.2333 | 0.2482 | $g$ |
| $\sigma_p$ | 280.6 | 372.3 | 531 | $\mu g.g^{-1}$ |
| $\sigma_{met1}$ | 42.07 | 56.26 | 80.95 | $\mu g.g^{-1}$ |
| $\sigma_{met2}$ | 103.5 | 140.1 | 207.8 | $\mu g.g^{-1}$ |
| $\sigma_{met3}$ | 34.28 | 46.33 | 68.97 | $\mu g.g^{-1}$ |
| $\sigma_G$ | 0.01243 | 0.01697 | 0.02507 | $g$ |

#### Goodness-of-fit criteria

##### Posterior Predictive Check

The PPC shows the observed values against their corresponding estimated predictions (black dots), along with their 95% credible interval (vertical segments). If the fit is correct, we expect to see 95% of the data within the intervals. Ideally observations and predictions should coincide, so we would expect to see black dots along the first bisector y = x (plain black line). The 95% credible intervals are colored in green if they overlap this line, in red otherwise.

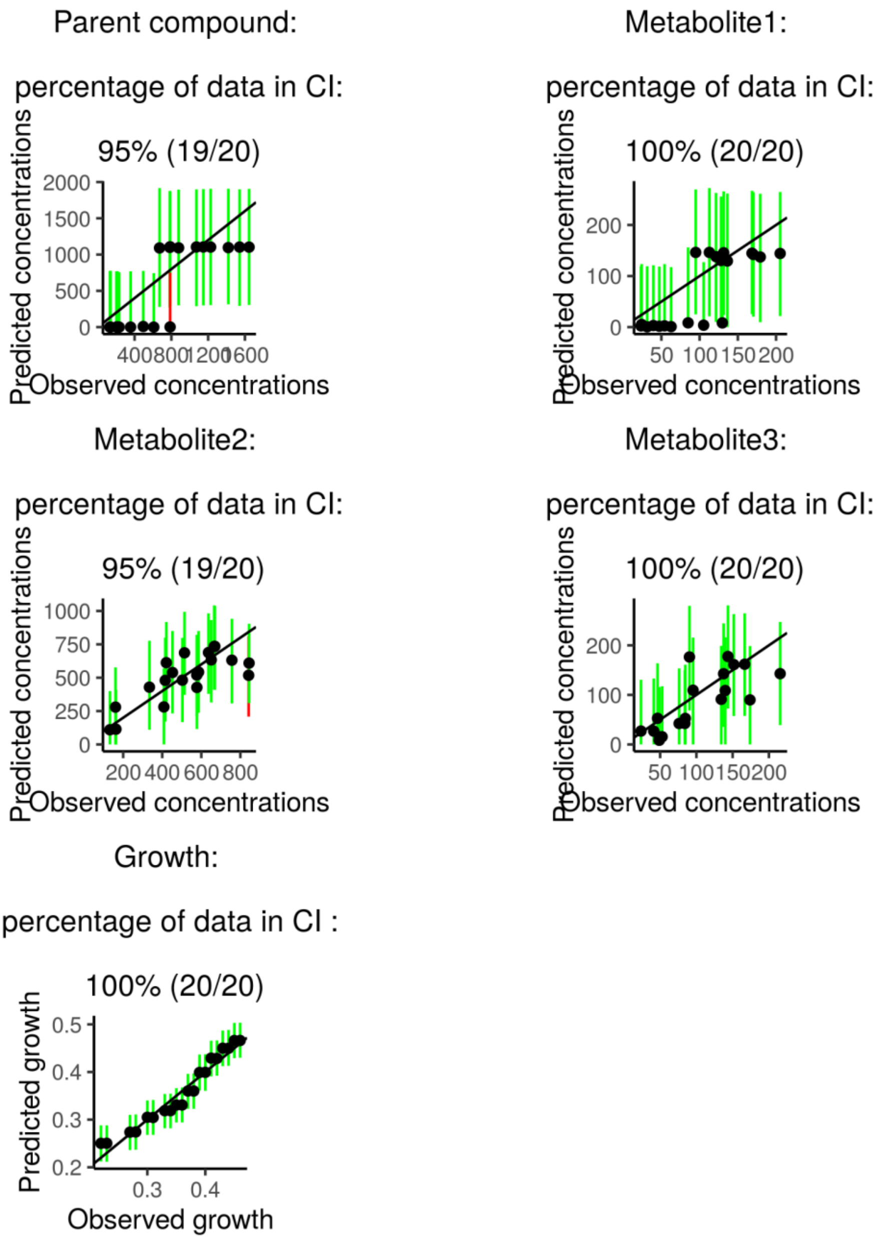

**Priors and posteriors**

The prior distribution is represented by the gray area and the posterior distribution by the orange area. The accuracy of the model parameter estimation can be visualized by comparing prior and posterior distributions: the overall expectation is to get a narrower posterior distribution compared to the prior one, what reflects that data contributed enough to precisely estimate parameters.

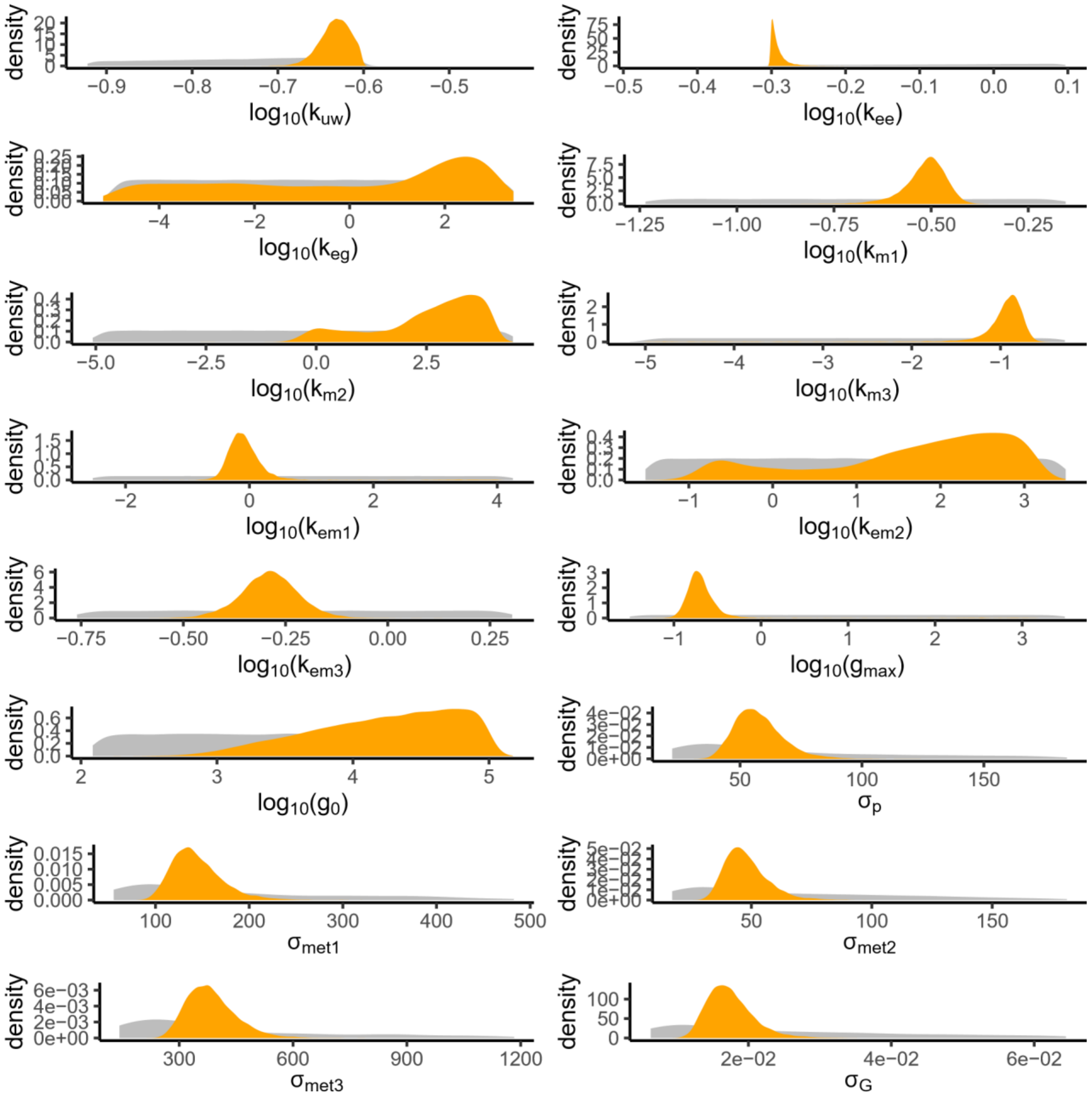

##### Correlation between parameters

Correlations between parameters are visualized by projecting the joint posterior distribution in a plot matrix with planes of parameter pairs (lower triangular elements), marginal posterior distribution of each model parameter (diagonal), and Pearson correlation coefficients (upper triangular elements). Correlations are expected to be low (reflected by “potatoid” shapes of density lines in orange); a leaning elliptical shape translates high correlations (positive if leaning to the right, negative if leaning to the left).

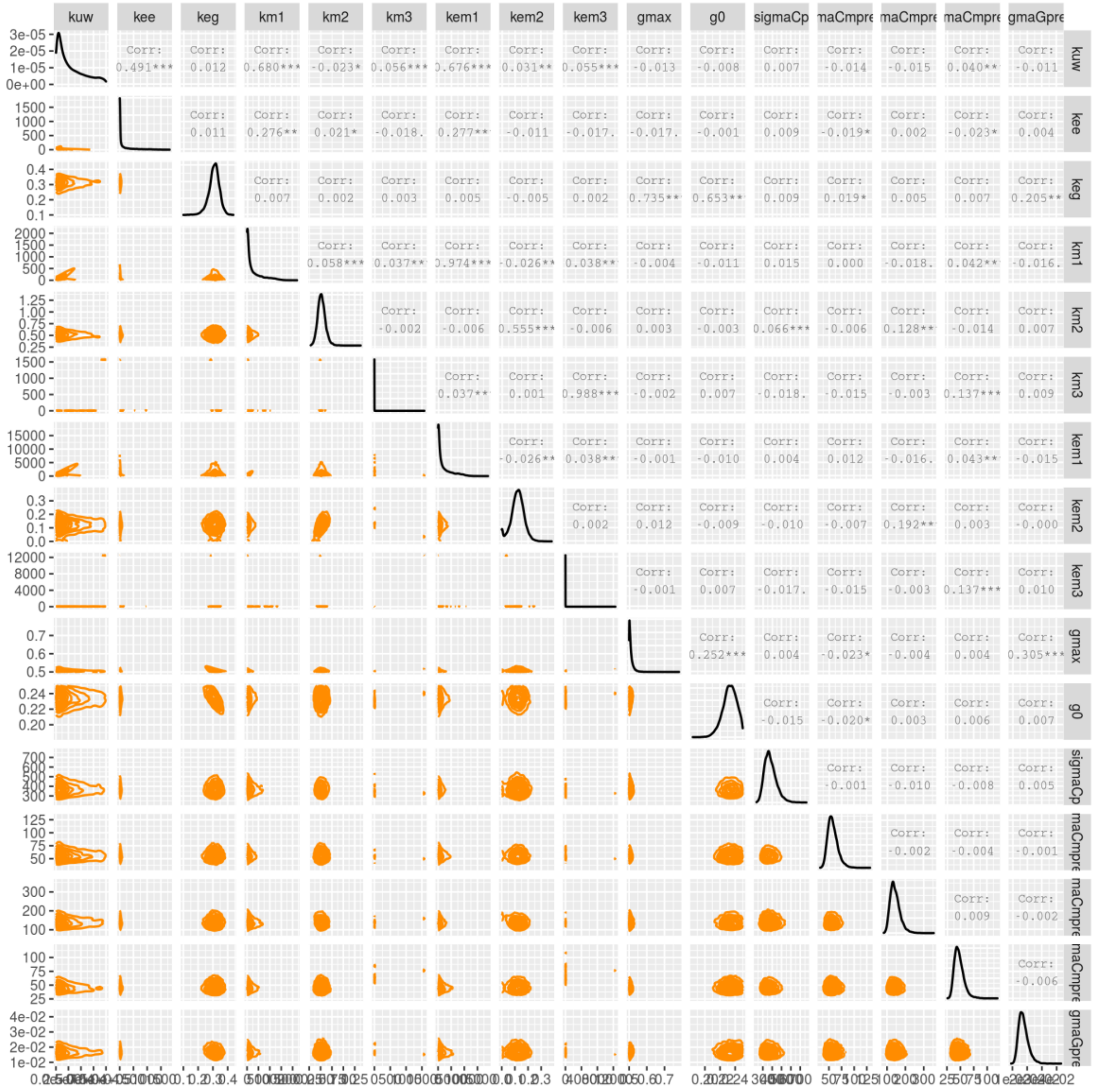

#### Color matrix of correlation between parameters

A colored matrix allows to see at a glance the most correlated (green) or anti-correlated (orange) parameters.

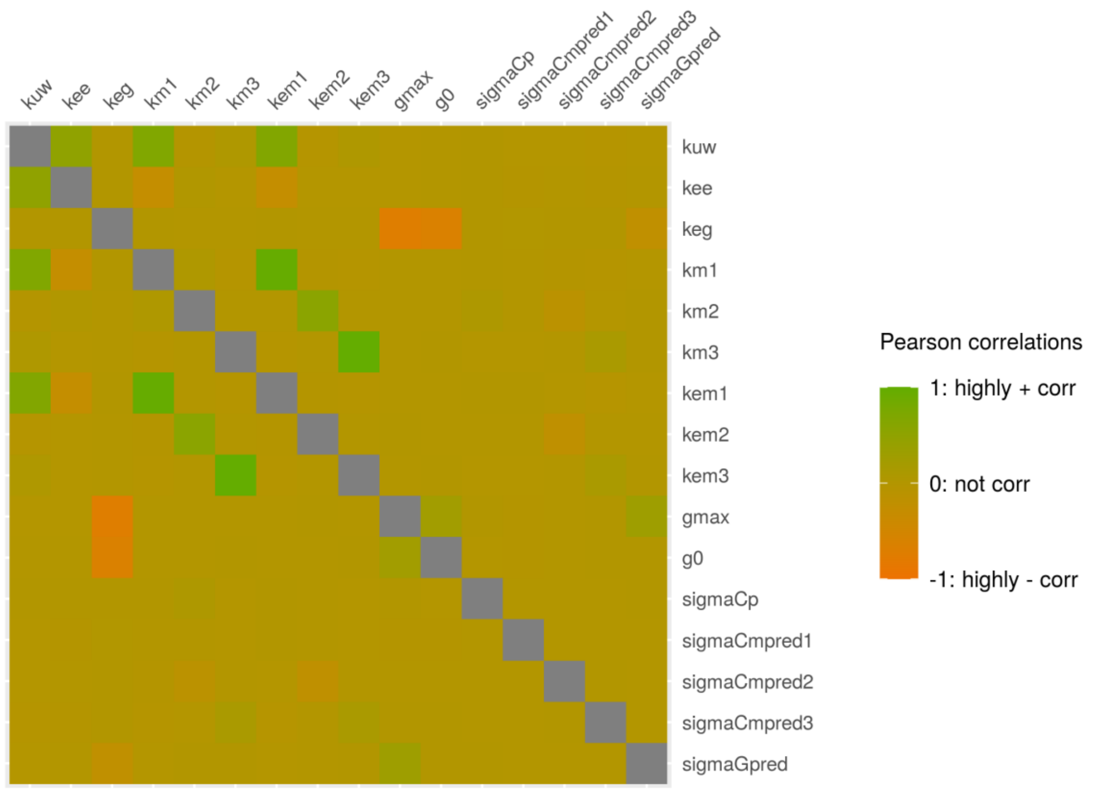

**Potential Scale Reduction Factors**

Convergence of the MCMC chains can be check with the Gelman-Rubin diagnostic expressed with the potential scale reduction factor (PSRF). Approximate convergence is diagnosed when the PSRF is close to 1.

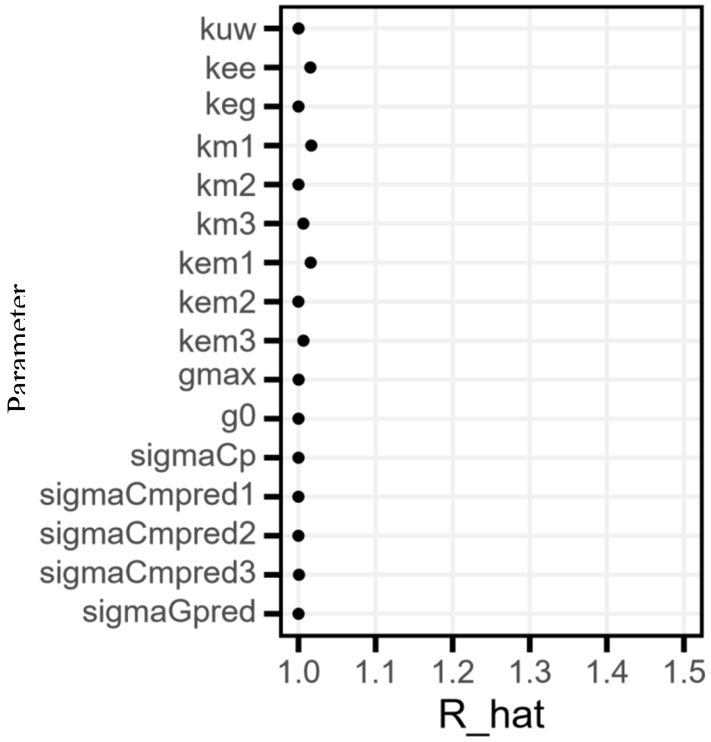


**Deviance Information Criterion**

This criteria, denoted DIC, is a penalized deviance statistics accounting for the number of parameters for use in model comparison for a same dataset (*e.g.*, with or without $k_e$). Sub-models with lower DIC values will be preferred.

**DIC = 873**

**Traces of MCMC iterations**

A traceplot is an essential plot for assessing convergence and diagnosing of MCMC chains. It shows the time series of the sampling process leading to the posterior distribution. Different colors are used for each of the chains (here 3) to assess within-chain convergence.

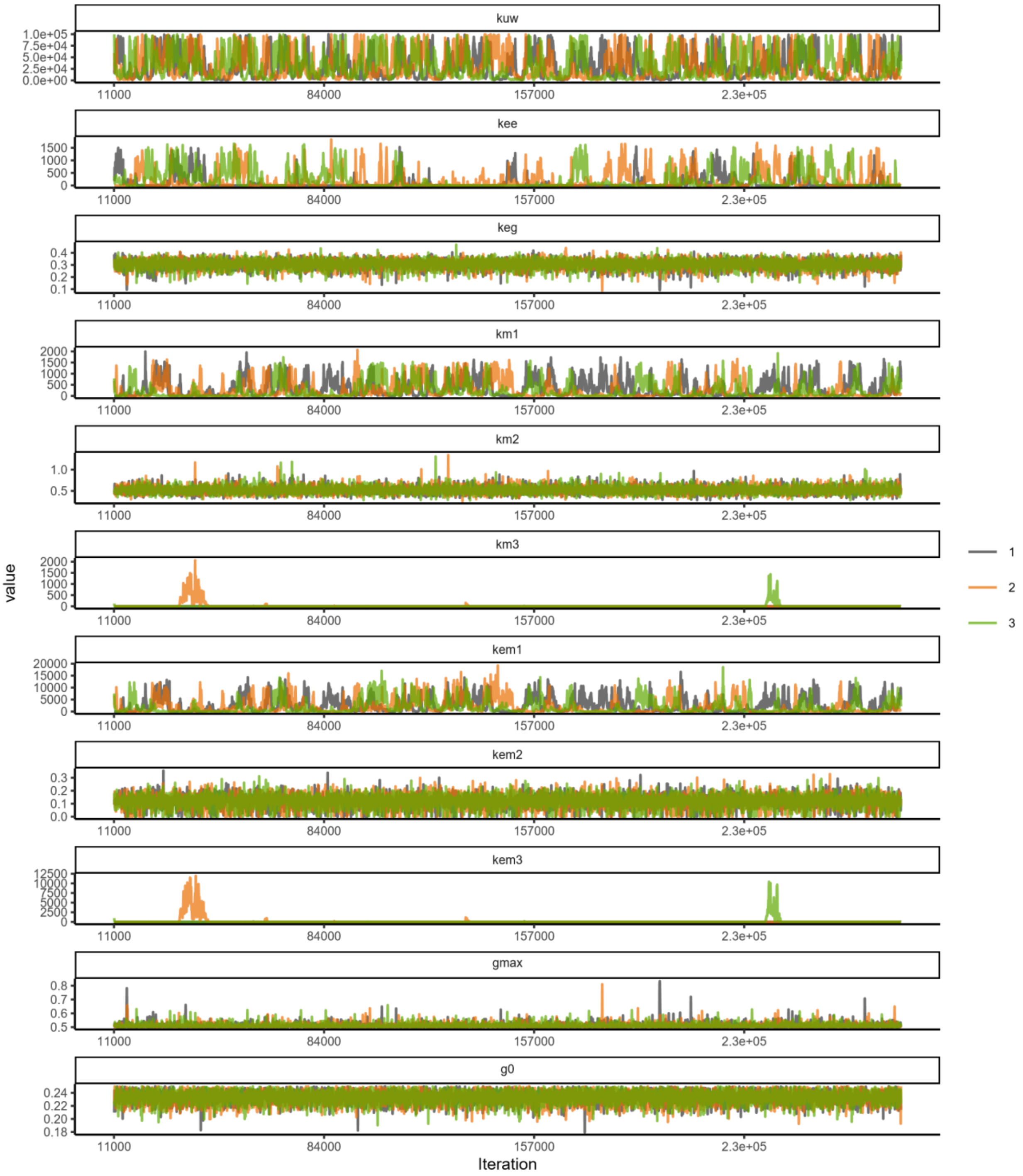

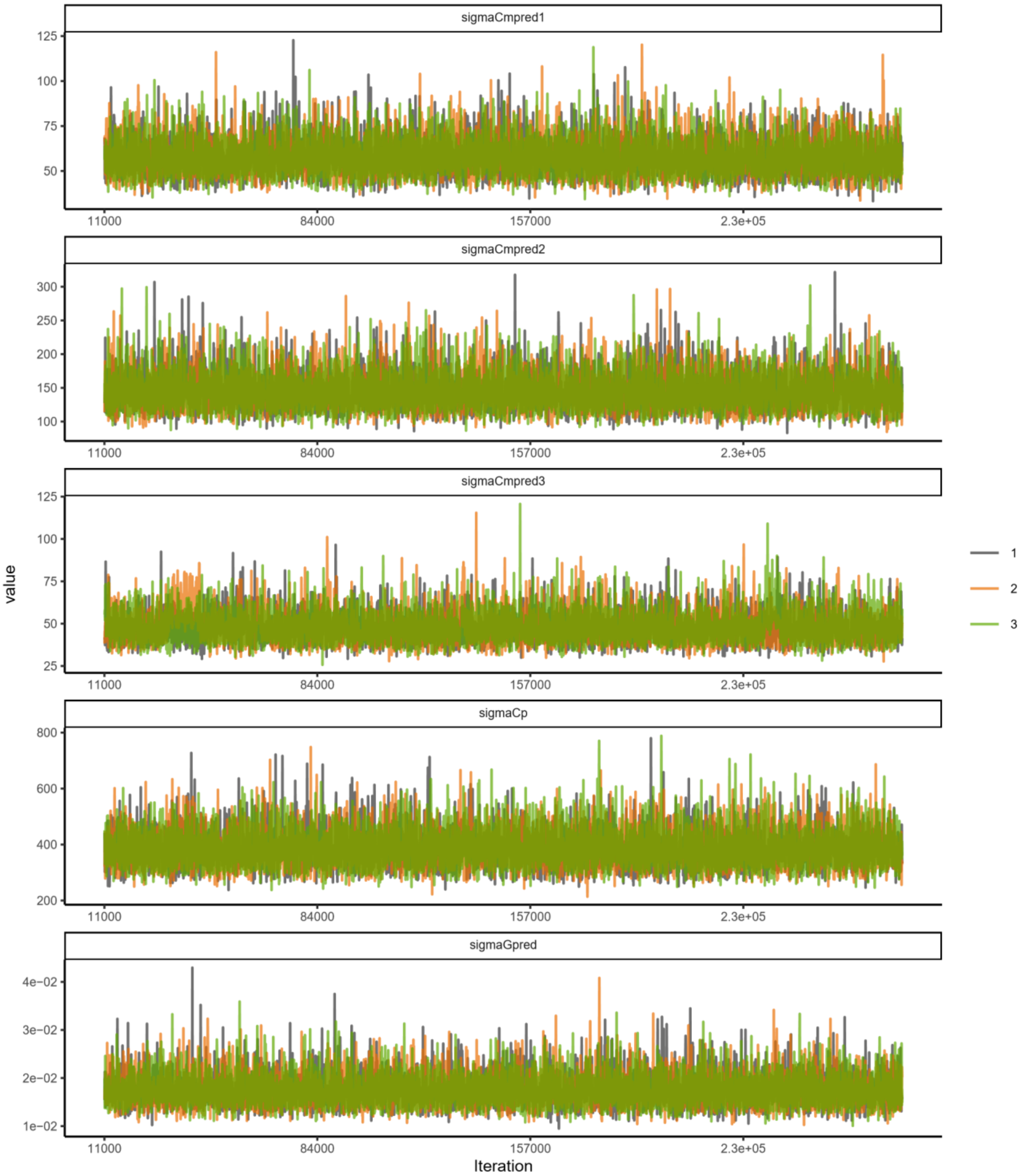
sigmaCmpred1
125
100
75
50
11000
84000
157000
2.3e+05
sigmaCmpred2
300
250
200
150
100
sigmaCmpred3
25
sigmaCp
800
600
400
200
sigmaGpred
4e−02
3e−02
2e−02
1e−02
value
Iteration
1
2
3

**Data Table**

Data used for the fitting process

| time | expw | conc | concm1 | concm2 | concm3 | replicate | growth |
|---|---|---|---|---|---|---|---|
| 0.000 | 15.53 | 0.0 | 0.00 | 0.0 | 0.00 | 1 | 0.20 |
| 0.000 | 15.53 | 0.0 | 0.00 | 0.0 | 0.00 | 2 | 0.20 |
| 0.207 | 15.53 | 878.6 | 128.07 | 132.1 | 84.06 | 1 | 0.23 |
| 0.208 | 15.53 | 782.8 | 136.32 | 162.9 | 76.18 | 2 | 0.22 |
| 0.520 | 15.53 | 1541.9 | 179.34 | 408.4 | 134.22 | 1 | 0.27 |
| 0.521 | 15.53 | 671.4 | 121.45 | 160.6 | 173.91 | 2 | 0.28 |
| 0.978 | 15.53 | 1645.1 | 205.36 | 842.4 | 137.51 | 1 | 0.31 |
| 0.979 | 15.53 | 1419.3 | 170.31 | 576.9 | 215.52 | 2 | 0.30 |
| 1.207 | 15.53 | 1227.1 | 131.51 | 756.6 | 166.57 | 1 | 0.33 |
| 1.208 | 15.53 | 1147.9 | 168.54 | 650.9 | 151.54 | 2 | 0.34 |
| 1.416 | 15.53 | 1072.5 | 94.87 | 669.2 | 143.49 | 1 | 0.35 |
| 1.417 | 15.53 | 786.4 | 112.74 | 666.1 | 90.42 | 2 | 0.36 |
| 1.999 | 15.53 | 494.1 | 84.94 | 636.7 | 95.41 | 1 | 0.37 |
| 2.000 | 15.53 | 786.3 | 129.75 | 514.0 | 139.77 | 2 | 0.38 |
| 2.978 | 15.53 | 608.5 | 105.30 | 844.5 | 84.69 | 1 | 0.39 |
| 2.979 | 15.53 | 232.0 | 24.12 | 421.0 | 46.39 | 2 | 0.40 |
| 3.999 | 15.53 | 222.6 | 54.13 | 586.0 | 41.18 | 1 | 0.41 |
| 4.000 | 15.53 | 356.4 | 39.53 | 452.4 | 23.57 | 2 | 0.42 |
| 4.999 | 15.53 | 213.1 | 47.30 | 503.0 | 48.84 | 1 | 0.43 |
| 5.000 | 15.53 | 131.3 | 23.32 | 415.5 | 52.70 | 2 | 0.44 |
| 5.999 | 15.53 | 201.9 | 62.72 | 577.4 | 48.85 | 1 | 0.45 |
| 6.000 | 15.53 | 138.6 | 31.38 | 334.4 | 48.51 | 2 | 0.46 |

#### ANNEX 4: EXAMPLE OF A REPORT PROVIDED BY MOSAIC$_{bioacc}$ WITH SEDIMENT EXPOSURE ROUTE

In order to illustrate MOSAIC$_{bioacc}$ with a sediment exposure route, the example file 'Chironomus_benzo-a-pyrene.csv' was selected in the application. In this example, male *Chironomus tentans* were exposed to benzo-(a)-pyrene spiked sediment and a single exposure concentration was tested. The duration of the accumulation phase is 3 days. One metabolite was quantified.

After calculations, the corresponding report is directly downloaded in MOSAIC$_{bioacc}$.

### MOSAIC$_{bioacc}$ REPORT

2021-04-19

This report is provided by the MOSAIC$_{bioacc}$ application available here: https://mosaic.univ-lyon1.fr/bioacc

Contact:

MOSAIC$_{bioacc}$ uses the JAGS (version 4.3.0) and R (version 4.0.2) software, and in particular packages RJags (version 4.10), jagsUI (version 1.5.1) and Shiny (version 1.6.0).

The MOSAIC$_{bioacc}$ application is a turn-key web tool providing bioaccumulation metrics (BCF/BSAF/BMF) from a toxicokinetic (TK) model fitted to accumulation-depuration data. It is designed to fulfil the requirements of regulators when examining applications for market authorization of active substances.

##### Data summary

File used: Chironomus_benzo-a-pyrene.csv

Exposure: 0.0136 $\mu g.g^{-1}$

Accumulation phase duration: 3 days

Number of replicates: 2

Times: 0, 0.25, 0.5, 0.75, 1, 1.5, 2, 3, 3.25, 3.5, 4, 4.625, 5, 6

Exposure routes: sediment

Elimination routes: excretion biotransformation

##### Bayesian inference

Three MCMC chains were used to estimate model parameters.

Number of iterations: 176062 Thin: 47

##### TK Model

The TK model used for these calculations was:

$$\frac{dC_p(t)}{dt} = k_{us} \times c_s - (k_{ee} + k_{m1}) \times C_p(t) \quad for\ 0 \le t \le t_c$$

$$\frac{dC_p(t)}{dt} = -(k_{ee} + k_{m1}) \times C_p(t) \quad for\ t > t_c$$

$$\frac{dC_{m1}(t)}{dt} = k_{m1} \times C_p(t) - k_{em1} \times C_{m1}(t)$$

with:

$t$: time (expressed in days )

$t_c$: duration of the accumulation phase (expressed in days )

$C_p(t)$: internal concentration of the parent compound at time (expressed in $\mu g.g^{-1}$)

$k_{ee}$: elimination rates of excretion (expressed per days $^{-1}$)

$c_s$: exposure concentration of sediment route (expressed in $\mu g.g^{-1}$)

$k_{us}$: uptake rate of sediment exposure (expressed per days $^{-1}$)

$C_{m\ell}(t)$: internal concentration of metabolite $\ell$ (expressed in $\mu g.g^{-1}$)

$\ell$: index of metabolites, $\ell = 1 \ldots L$ with $L$ total number of metabolites

$k_{m\ell}$: metabolization rate of metabolite $\ell$ (expressed per days $^{-1}$)

$k_{em\ell}$: elimination rates of metabolite $\ell$ (expressed per days $^{-1}$)

#### Bioaccumulation metric calculation

##### Calculations

$$BSAF_k = \frac{k_{us}}{k_{ee} + k_{m1}}$$

$$BSAF_{ss} = \frac{C_p(t_c)}{c_s}$$

#### Biote-sediment accumulation factor (BSAF)

##### $BSAF_k$ plot

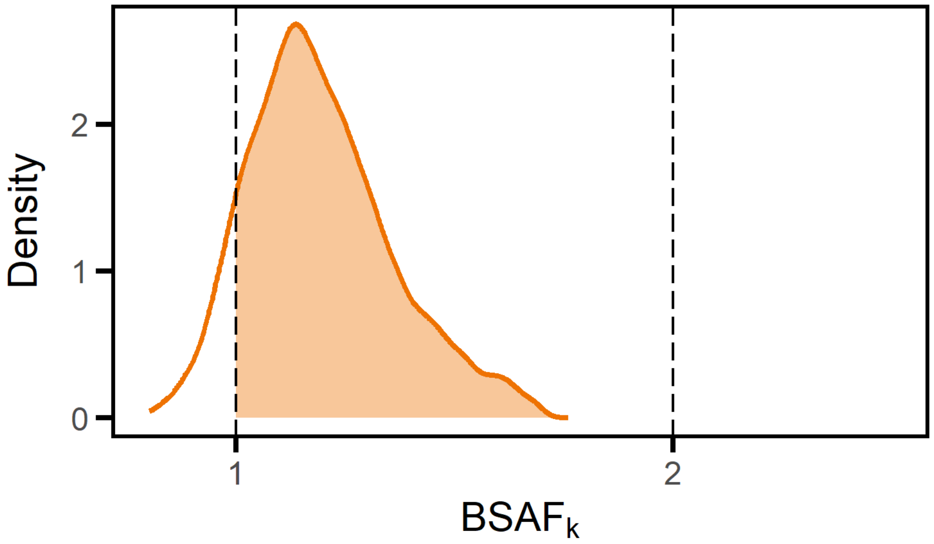


##### $BSAF_{ss}$ plot

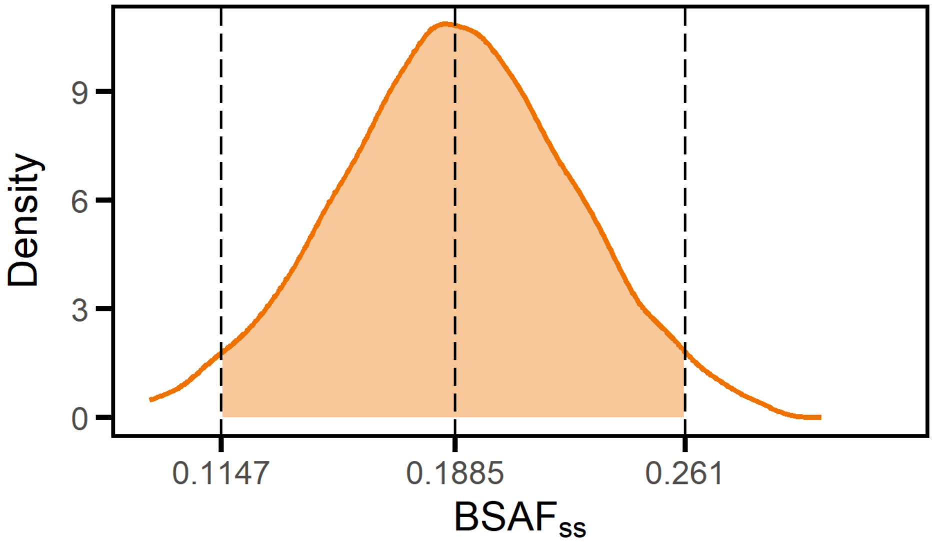


##### BSAF summary

| | 2.5% | 50% | 97.5% | CV |
|---|---|---|---|---|
| BSAFk | 1 | 1 | 2 | 0.25 |
| BSAFss | 0.1147 | 0.1885 | 0.261 | 0.19 |

**Fitting results**

**Fit plot**

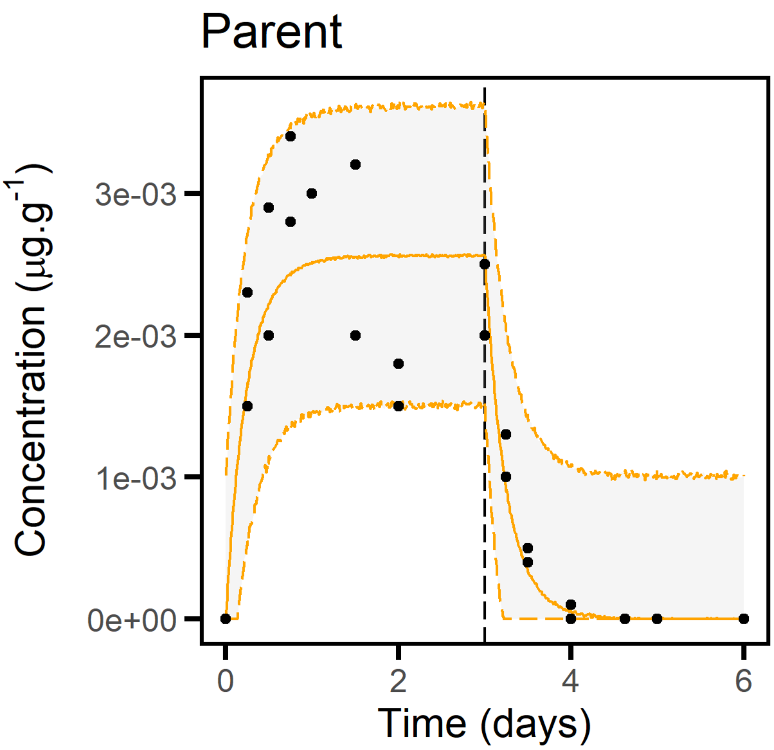


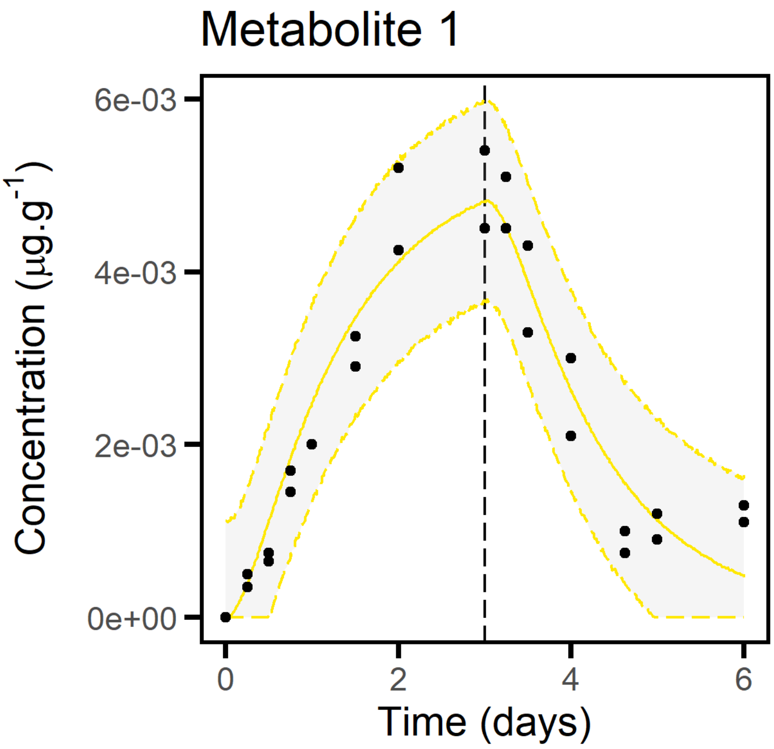

##### Quantiles of estimated parameters

| | 2.5% | 50% | 97.5% | |
|---|---|---|---|---|
| $k_{us}$ | 0.5732 | 149.2 | 12280 | $d^{-1}$ |
| $k_{ee}$ | 0.9983 | 846 | 70280 | $d^{-1}$ |
| $k_{m1}$ | 1.164 | 1.535 | 2.149 | $d^{-1}$ |
| $k_{em1}$ | 0.4781 | 0.6391 | 1.035 | $d^{-1}$ |
| $\sigma_p$ | 3.98e-04 | 5.66e-04 | 8.032e-04 | $\mu g.g^{-1}$ |
| $\sigma_{met1}$ | 4.367e-04 | 6.198e-04 | 8.91e-04 | $\mu g.g^{-1}$ |

**Goodness-of-fit criteria**

##### Posterior Predictive Check

The PPC shows the observed values against their corresponding estimated predictions (black dots), along with their 95% credible interval (vertical segments). If the fit is correct, we expect to see 95% of the data within the intervals. Ideally observations and predictions should coincide, so we would expect to see black dots along the first bisector y = x (plain black line). The 95% credible intervals are colored in green if they overlap this line, in red otherwise.

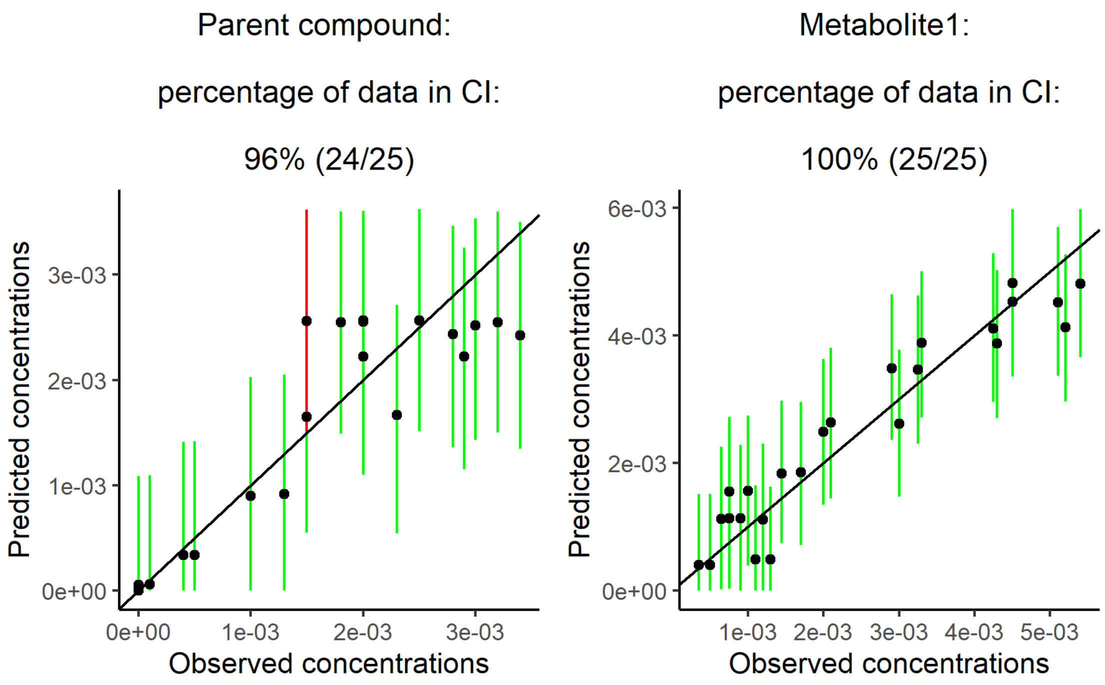


##### Priors and posteriors

The prior distribution is represented by the gray area and the posterior distribution by the orange area. The accuracy of the model parameter estimation can be visualized by comparing prior and posterior distributions: the overall expectation is to get a narrower posterior distribution compared to the prior one, what reflects that data contributed enough to precisely estimate parameters.

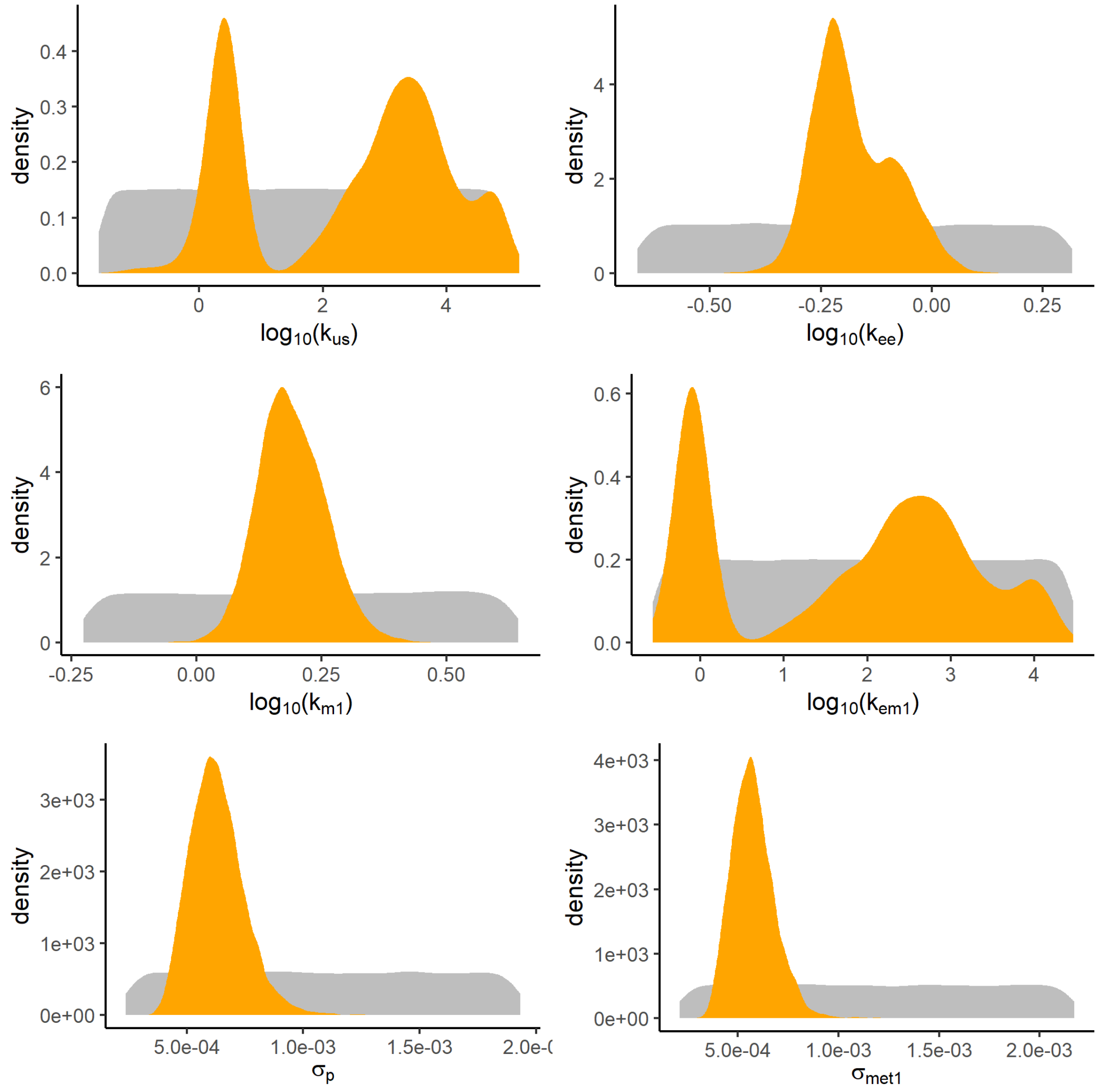


#### Color matrix of correlation between parameters

A colored matrix allows to see at a glance the most correlated (green) or anti-correlated (orange) parameters.

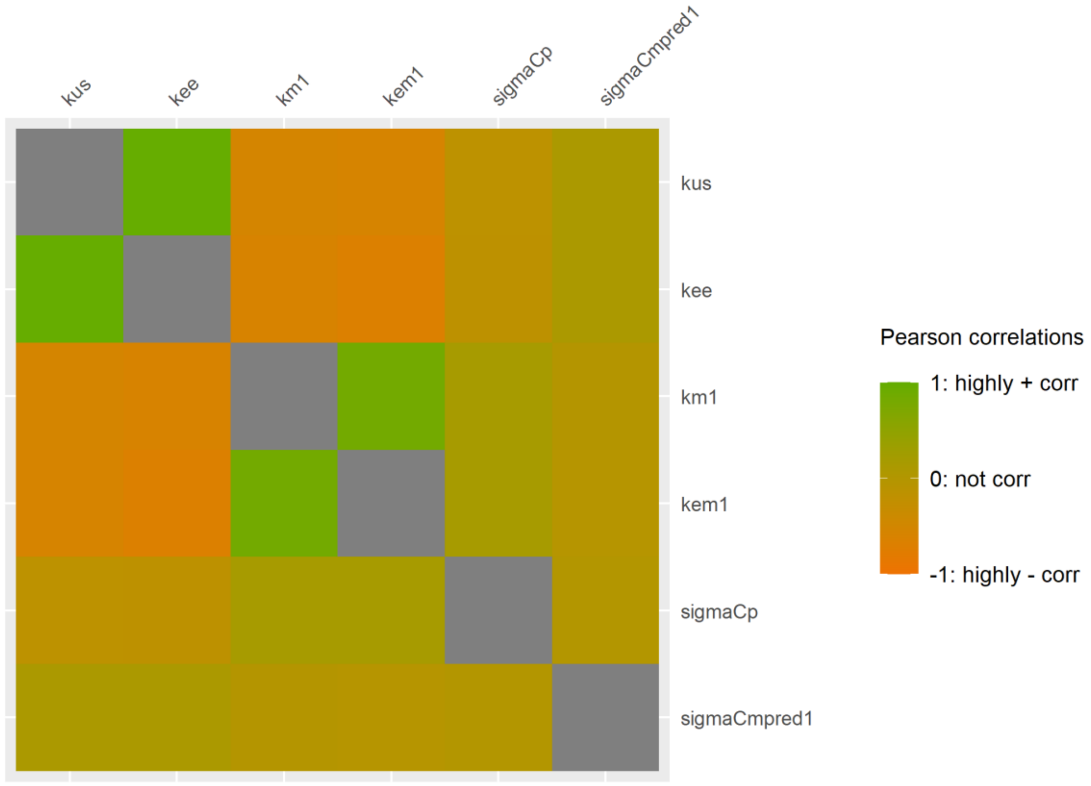


**Correlation between parameters**

Correlations between parameters are visualized by projecting the joint posterior distribution in a plot matrix with planes of parameter pairs (lower triangular elements), marginal posterior distribution of each model parameter (diagonal), and Pearson correlation coefficients (upper triangular elements). Correlations are expected to be low (reflected by “potatoid” shapes of density lines in orange); a leaning elliptical shape translates high correlations (positive if leaning to the right, negative if leaning to the left).

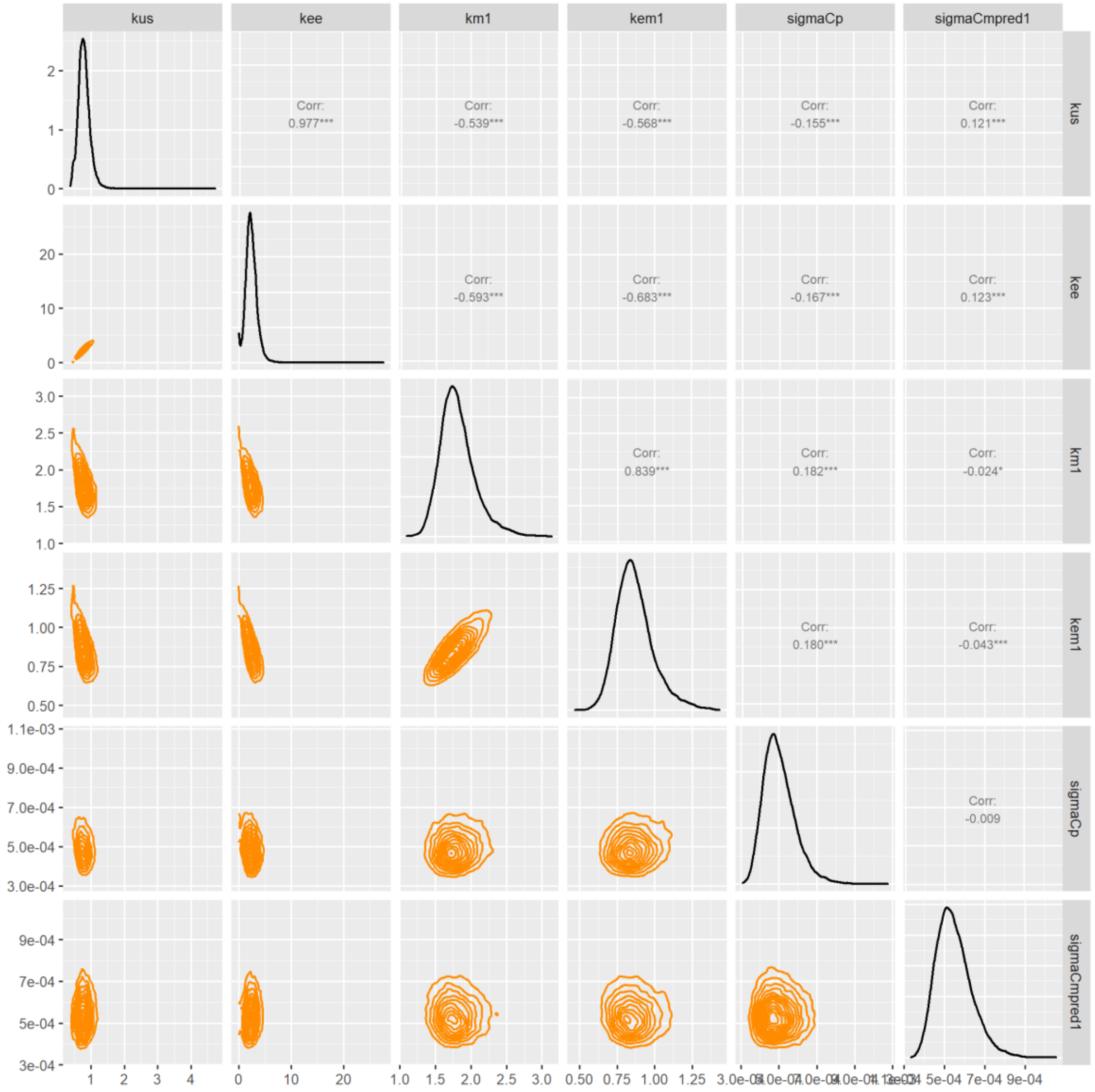


#### Potential Scale Reduction Factors

Convergence of the MCMC chains can be check with the Gelman-Rubin diagnostic expressed with the potential scale reduction factor (PSRF). Approximate convergence is diagnosed when the PSRF is below 1.01.

| | PSRF |
|---|---|
| $k_{us}$ | 1.022 |
| $k_{ee}$ | 1.017 |
| $k_{m1}$ | 1.003 |
| $k_{em1}$ | 1.003 |
| $\sigma_p$ | 1.000 |
| $\sigma_{met1}$ | 1.000 |

##### Watanabe–Akaike information criterion

Information criteria offer a computationally appealing way of estimating the generalization performance of the model. A fully Bayesian criterion is the widely applicable information criterion (WAIC) by Watanabe a penalized deviance statistics accounting for the uncertainty in the parameters and can be used also for singular models. WAIC is widely used in model comparison for a same dataset (e.g., with or without $k_{ee}$). Sub-models with lower WAIC values will be preferred.

WAIC = -609.9

##### Deviance Information Criterion

This criteria, denoted DIC, is a penalized deviance statistics accounting for the number of parameters for use in model comparison for a same dataset (e.g., with or without $k_{ee}$). Sub-models with lower DIC values will be preferred.

DIC = -607

##### Traces of MCMC iterations

A traceplot is an essential plot for assessing convergence and diagnosing of MCMC chains. It shows the time series of the sampling process leading to the posterior distribution. Different colors are used for each of the chains (here 3) to assess within-chain convergence.

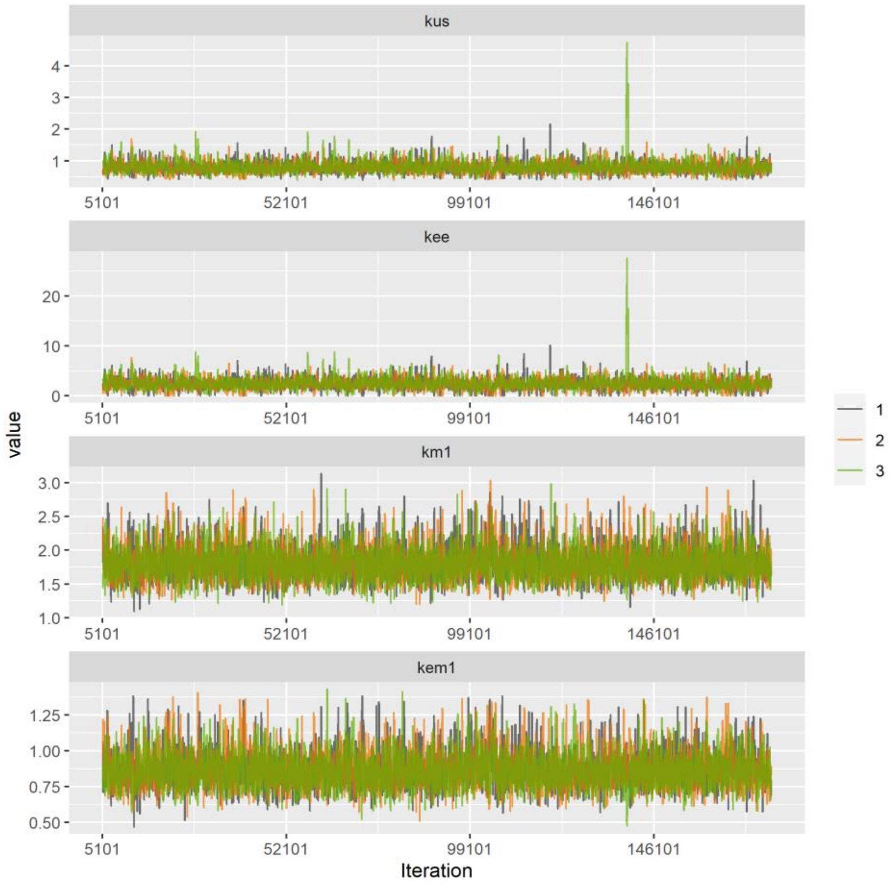

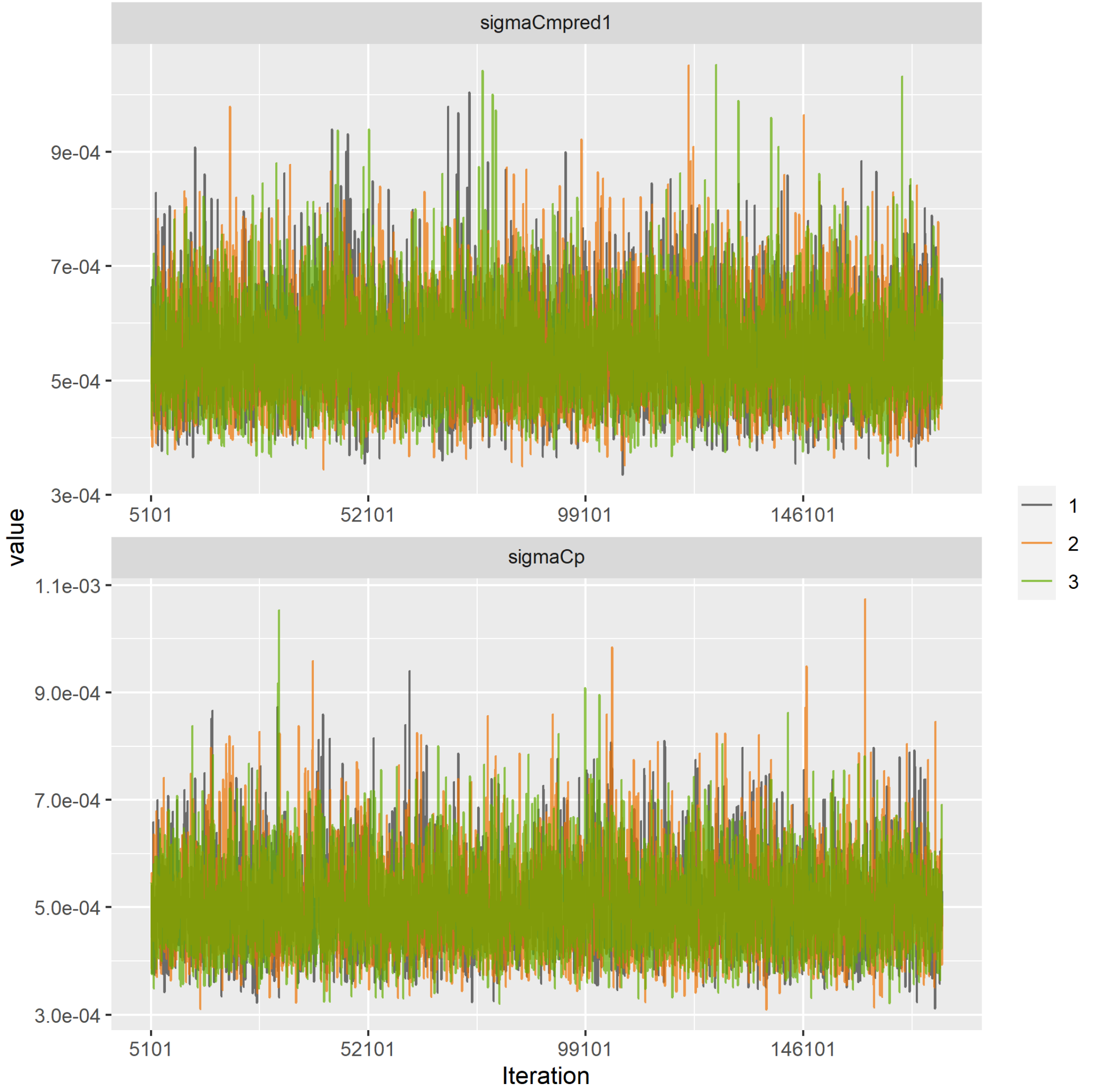
sigmaCmpred1
9e-04
7e-04
5e-04
3e-04
5101
52101
99101
146101
value
sigmaCp
1.1e-03
9.0e-04
7.0e-04
5.0e-04
3.0e-04
Iteration
1
2
3

#### Data Table

| time | exps | conc | concm1 | replicate |
|---|---|---|---|---|
| 0.000 | 0.0136 | 0.0000 | 0.0000 | 1 |
| 0.000 | 0.0136 | 0.0000 | 0.0000 | 2 |
| 0.250 | 0.0136 | 0.0015 | 0.0005 | 1 |
| 0.250 | 0.0136 | 0.0023 | 0.0004 | 2 |
| 0.500 | 0.0136 | 0.0020 | 0.0008 | 1 |
| 0.500 | 0.0136 | 0.0029 | 0.0006 | 2 |
| 0.750 | 0.0136 | 0.0028 | 0.0014 | 1 |
| 0.750 | 0.0136 | 0.0034 | 0.0017 | 2 |
| 1.000 | 0.0136 | 0.0030 | 0.0020 | 1 |
| 1.500 | 0.0136 | 0.0032 | 0.0029 | 1 |
| 1.500 | 0.0136 | 0.0020 | 0.0032 | 2 |
| 2.000 | 0.0136 | 0.0015 | 0.0043 | 1 |
| 2.000 | 0.0136 | 0.0018 | 0.0052 | 2 |
| 3.000 | 0.0136 | 0.0025 | 0.0045 | 1 |
| 3.000 | 0.0136 | 0.0020 | 0.0054 | 2 |
| 3.250 | 0.0136 | 0.0010 | 0.0045 | 1 |
| 3.250 | 0.0136 | 0.0013 | 0.0051 | 2 |
| 3.500 | 0.0136 | 0.0005 | 0.0043 | 1 |
| 3.500 | 0.0136 | 0.0004 | 0.0033 | 2 |
| 4.000 | 0.0136 | 0.0001 | 0.0021 | 1 |
| 4.000 | 0.0136 | 0.0000 | 0.0030 | 2 |
| 4.625 | 0.0136 | 0.0000 | 0.0010 | 1 |
| 4.625 | 0.0136 | 0.0000 | 0.0008 | 2 |
| 5.000 | 0.0136 | 0.0000 | 0.0009 | 1 |
| 5.000 | 0.0136 | 0.0000 | 0.0012 | 2 |
| 6.000 | 0.0136 | 0.0000 | 0.0011 | 1 |
| 6.000 | 0.0136 | 0.0000 | 0.0013 | 2 |

**ANNEX 5: EXAMPLE OF A REPORT PROVIDED BY MOSAIC$_{bioacc}$ WITH FOOD EXPOSURE ROUTE**

In order to illustrate MOSAIC$_{bioacc}$ with a food exposure route, the file' Solea_PCB153_90d_Eichinger2010.csv' was selected in the database and then uploaded in the application. In this example, male *Solea solea* were exposed to PCB153 spiked food and a single exposure concentration was tested. The duration of the accumulation phase is 90 days.

After calculations, the corresponding report is directly downloaded in MOSAIC$_{bioacc}$.

### MOSAIC$_{bioacc}$ REPORT

2021-04-19

---

This report is provided by the MOSAIC$_{bioacc}$ application available here: https://mosaic.univ-lyon1.fr/bioacc

Contact:

MOSAIC$_{bioacc}$ uses the JAGS (version 4.3.0) and R (version 4.0.2) software, and in particular packages RJags (version 4.10), jagsUI (version 1.5.1) and Shiny (version 1.6.0).

The MOSAIC$_{bioacc}$ application is a turn-key web tool providing bioaccumulation metrics (BCF/BSAF/BMF) from a toxicokinetic (TK) model fitted to accumulation-depuration data. It is designed to fulfil the requirements of regulators when examining applications for market authorization of active substances.

---

##### Data summary

File used: Solea_PCB153_90d_Eichinger2010.csv

Exposure: 888 $\mu g.g^{-1}$

Accumulation phase duration: 90 days

Number of replicates: 3

Times: 0, 4, 8, 14, 28, 56, 84, 88, 90, 98, 112, 140, 168

Exposure routes: food

Elimination routes: excretion

##### Bayesian inference

Three MCMC chains were used to estimate model parameters.

Number of iterations: 89904 Thin: 24

##### TK Model

The TK model used for these calculations was:

$$\frac{dC_p(t)}{dt} = k_{uf} \times c_f - (k_{ee}) \times C_p(t) \quad for\ 0 \le t \le t_c$$

$$\frac{dC_p(t)}{dt} = -(k_{ee}) \times C_p(t) \quad for\ t > t_c$$

with:

$t$: time (expressed in days )

$t_c$: duration of the accumulation phase (expressed in days )

$C_p(t)$: internal concentration of the parent compound at time (expressed in $\mu g. g^{-1}$)

$k_{ee}$: elimination rates of excretion (expressed per days $^{-1}$)

$c_f$: exposure concentration of food route (expressed in $\mu g. g^{-1}$)

$k_{uf}$: uptake rate of food exposure (expressed per days $^{-1}$)

**Bioaccumulation metric calculation**

**Calculations**

$$BMF_k = \frac{k_{uf}}{k_{ee}}$$

**$BMF_k$ plot**

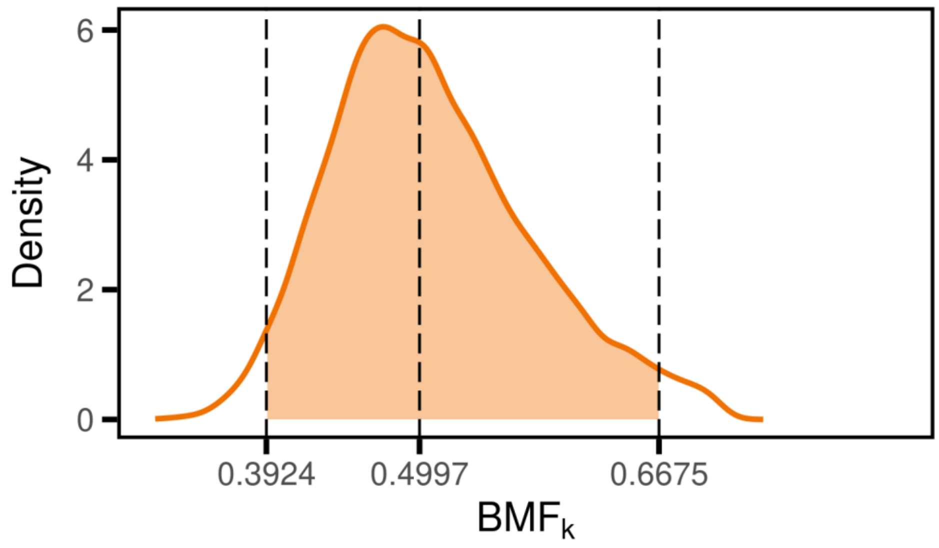


You didn’t ask for the $BMF_{ss}$

**BMF summary**

| | 2.5% | 50% | 97.5% | CV |
|---|---|---|---|---|
| BMFk | 0.3924 | 0.4997 | 0.6675 | 0.14 |

**Fitting results**

**Fit plot**

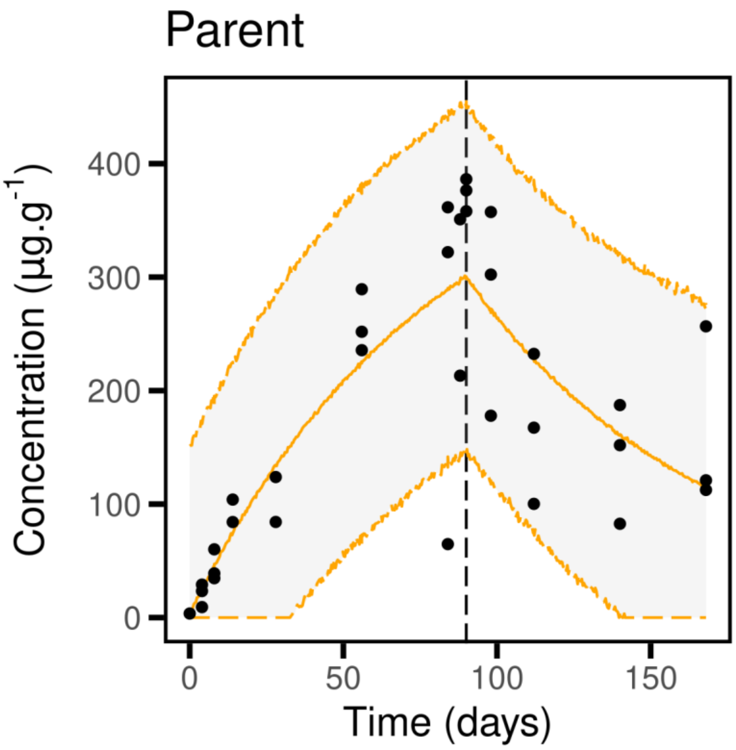


**Quantiles of estimated parameters**

| | 2.5% | 50% | 97.5% | |
|---|---|---|---|---|
| $k_{uf}$ | 4.285e-03 | 6.265e-03 | 8.644e-03 | $d^{-1}$ |
| $k_{ee}$ | 5.467e-03 | 0.01249 | 0.02046 | $d^{-1}$ |
| $\sigma_p$ | 57.82 | 73.53 | 98.42 | $\mu g.g^{-1}$ |

**Goodness-of-fit criteria**

**Posterior Predictive Check**

The PPC shows the observed values against their corresponding estimated predictions (black dots), along with their 95% credible interval (vertical segments). If the fit is correct, we expect to see 95% of the data within the intervals. Ideally observations and predictions should coincide, so we would expect to see black dots along the first bisector y = x (plain black line). The 95% credible intervals are colored in green if they overlap this line, in red otherwise.

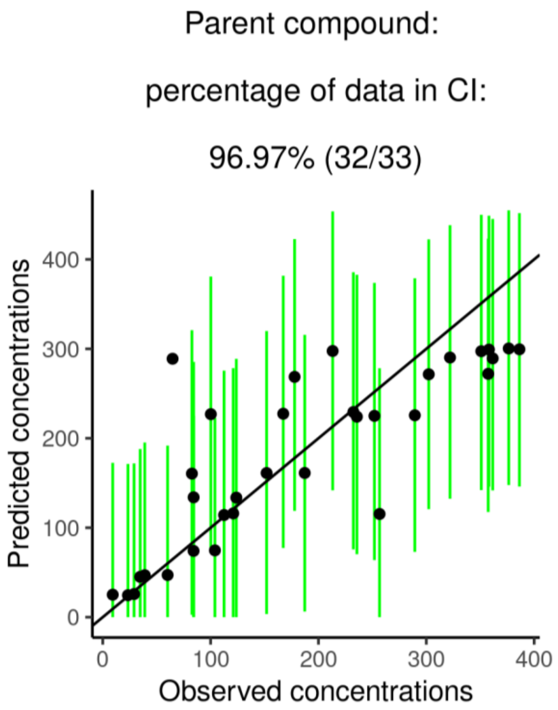


**Priors and posteriors**

The prior distribution is represented by the gray area and the posterior distribution by the orange area. The accuracy of the model parameter estimation can be visualized by comparing prior and posterior distributions: the overall expectation is to get a narrower posterior distribution compared to the prior one, what reflects that data contributed enough to precisely estimate parameters.

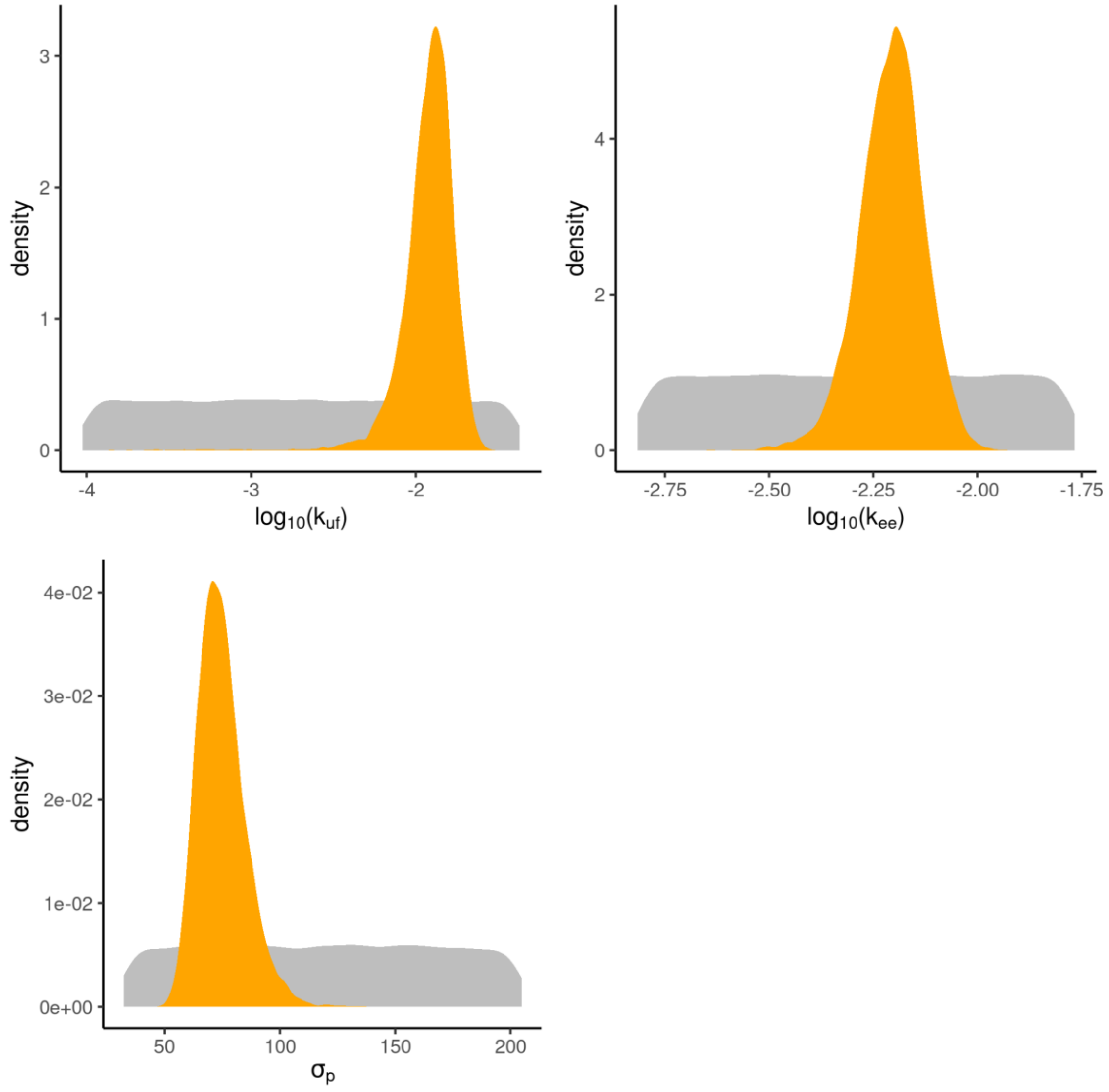


#### Color matrix of correlation between parameters

A colored matrix allows to see at a glance the most correlated (green) or anti-correlated (orange) parameters.

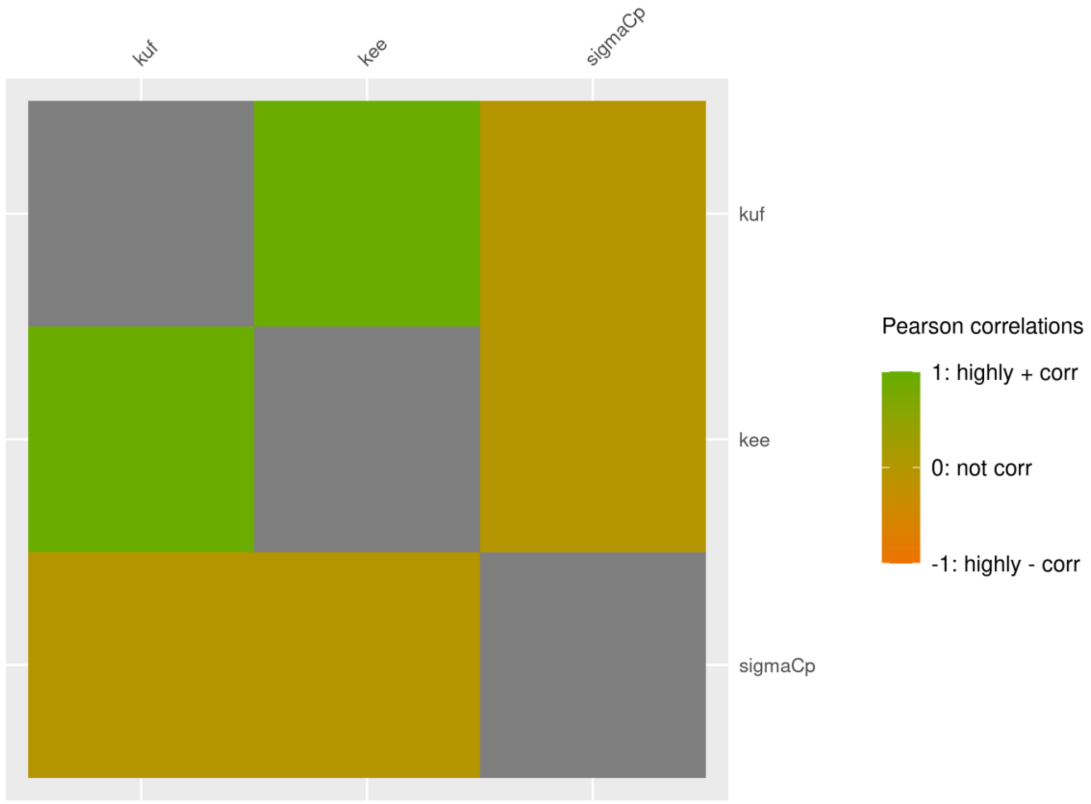


#### Correlation between parameters

Correlations between parameters are visualized by projecting the joint posterior distribution in a plot matrix with planes of parameter pairs (lower triangular elements), marginal posterior distribution of each model parameter (diagonal), and Pearson correlation coefficients (upper triangular elements). Correlations are expected to be low (reflected by “potatoid” shapes of density lines in orange); a leaning elliptical shape translates high correlations (positive if leaning to the right, negative if leaning to the left).

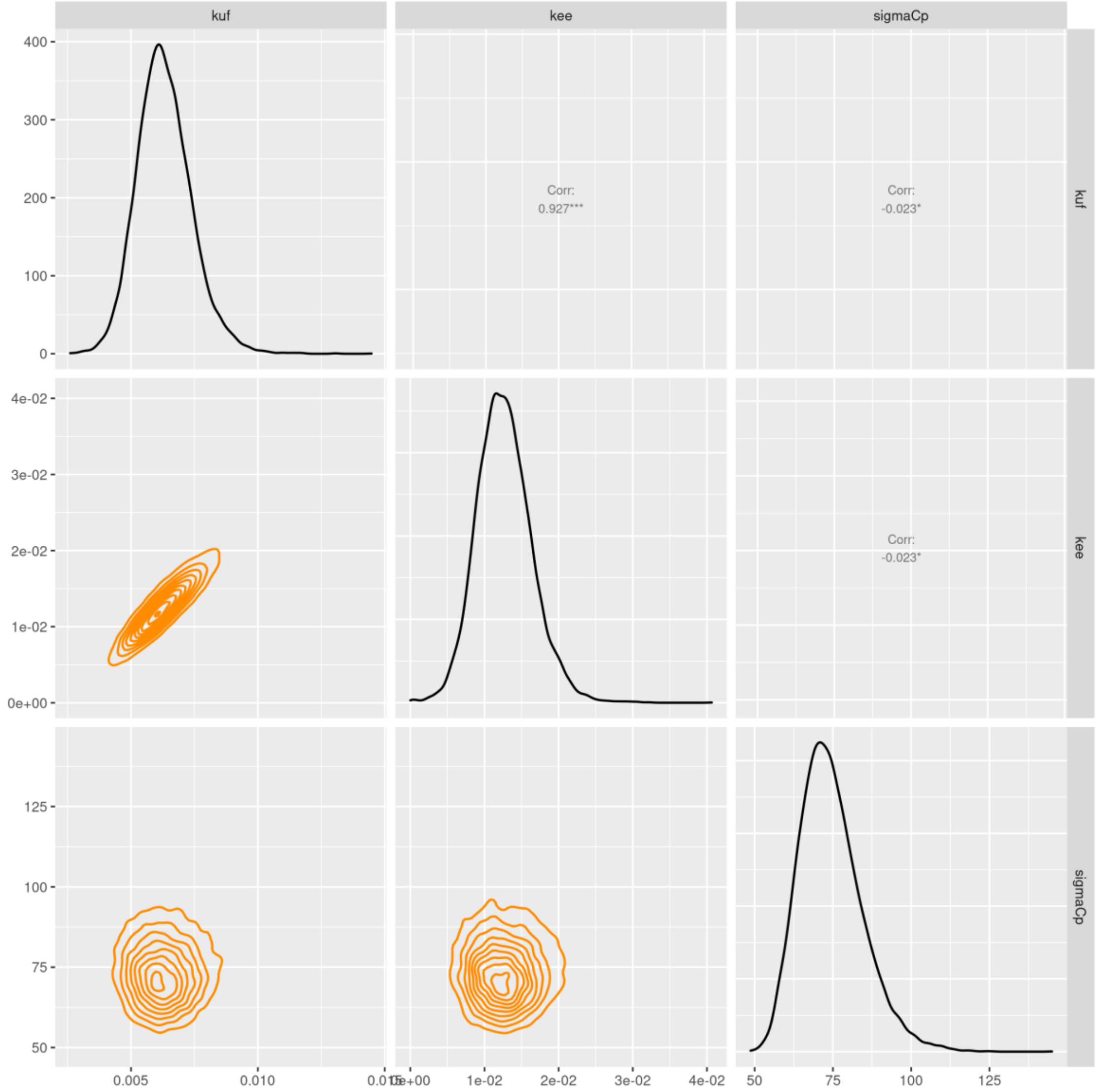


**Potential Scale Reduction Factors**

Convergence of the MCMC chains can be check with the Gelman-Rubin diagnostic expressed with the potential scale reduction factor (PSRF). Approximate convergence is diagnosed when the PSRF is below 1.01.

| | PSRF |
|---|---|
| $k_{uf}$ | 1.000 |
| $k_{ee}$ | 1.000 |
| $\sigma_p$ | 1.000 |

**Watanabe–Akaike information criterion**

Information criteria offer a computationally appealing way of estimating the generalization performance of the model. A fully Bayesian criterion is the widely applicable information criterion (WAIC) by Watanabe a penalized deviance statistics accounting for the uncertainty in the parameters and can be used also for singular models. WAIC is widely used in model comparison for a same dataset (e.g., with or without $k_{ee}$). Sub-models with lower WAIC values will be preferred.

WAIC = 380.9

##### Deviance Information Criterion

This criteria, denoted DIC, is a penalized deviance statistics accounting for the number of parameters for use in model comparison for a same dataset (e.g., with or without $k_{ee}$). Sub-models with lower DIC values will be preferred.

DIC = 380.5

##### Traces of MCMC iterations

A traceplot is an essential plot for assessing convergence and diagnosing of MCMC chains. It shows the time series of the sampling process leading to the posterior distribution. Different colors are used for each of the chains (here 3) to assess within-chain convergence.

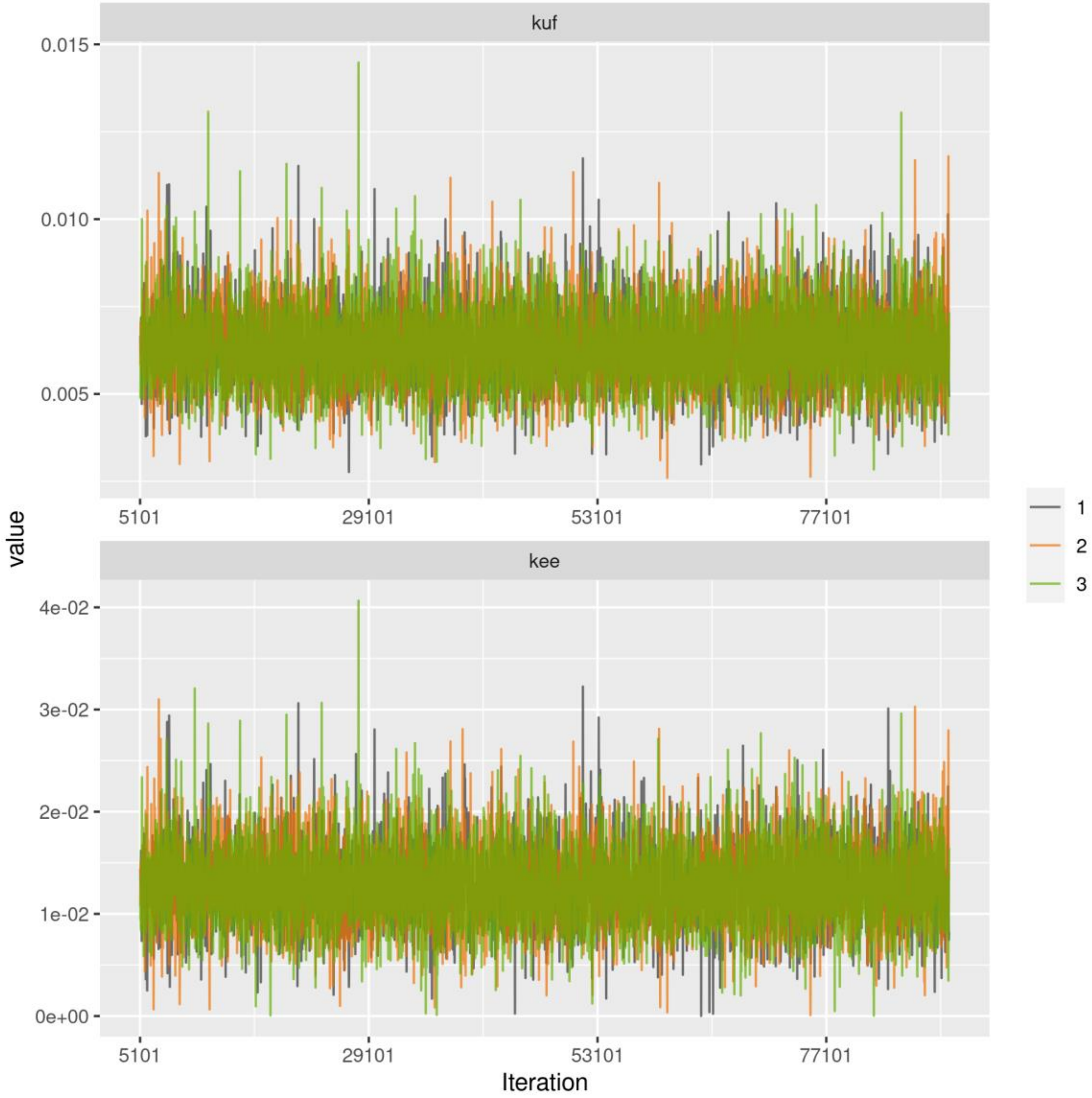
kuf
0.015
0.010
0.005
5101
29101
53101
77101
value
kee
4e-02
3e-02
2e-02
1e-02
0e+00
5101
29101
53101
77101
Iteration
1
2
3

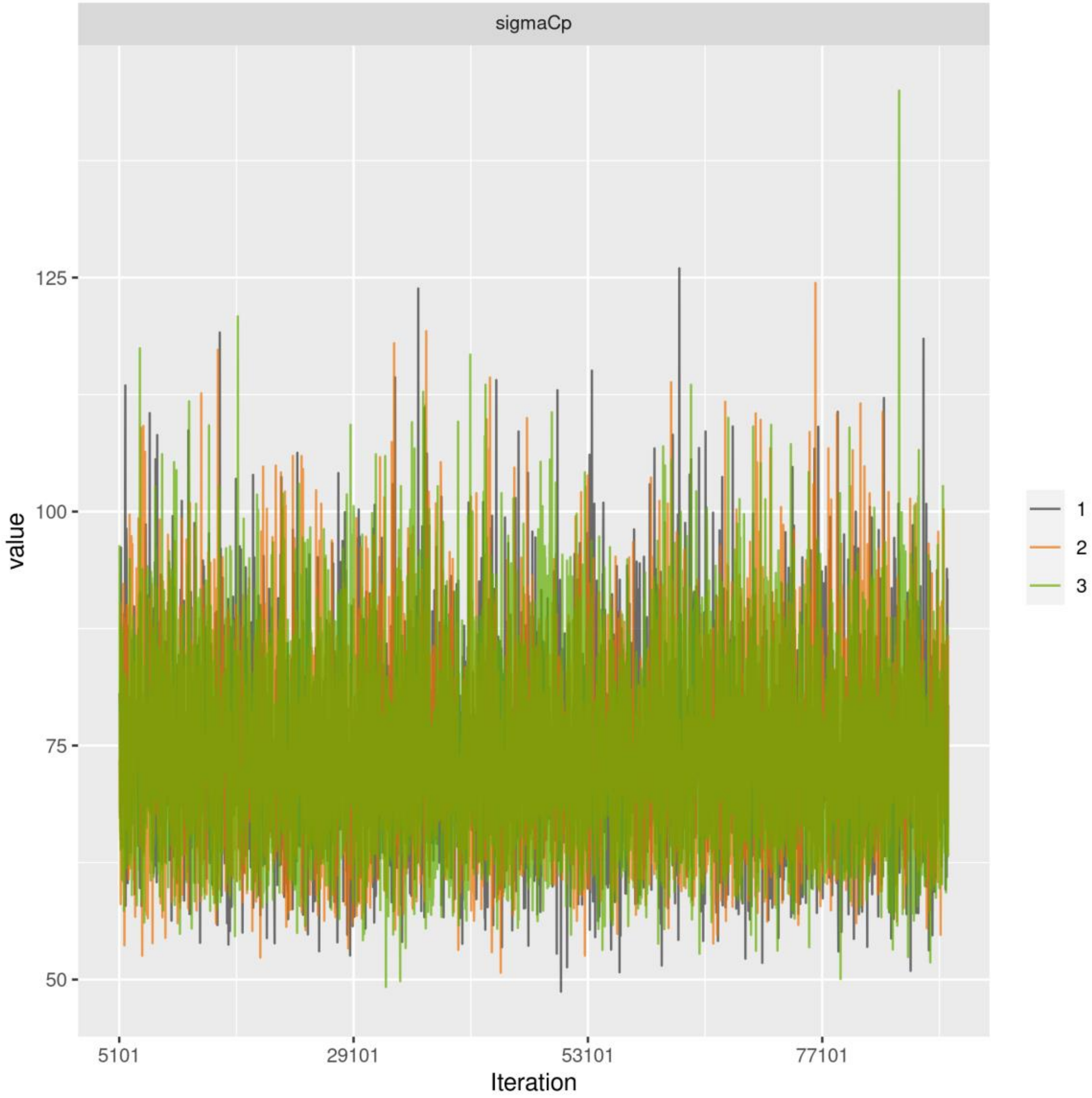
sigmaCp
125
100
75
50
value
5101
29101
53101
77101
Iteration
1
2
3

**Data Table**

| time | conc | expf | replicate |
|---|---|---|---|
| 0 | 3.566 | 888 | 1 |
| 4 | 9.237 | 888 | 1 |
| 4 | 23.372 | 888 | 2 |
| 4 | 29.026 | 888 | 3 |
| 8 | 34.700 | 888 | 1 |
| 8 | 38.940 | 888 | 2 |
| 8 | 60.141 | 888 | 3 |
| 14 | 84.197 | 888 | 1 |
| 14 | 103.985 | 888 | 2 |
| 28 | 84.263 | 888 | 1 |
| 28 | 123.839 | 888 | 2 |
| 56 | 235.632 | 888 | 1 |
| 56 | 251.886 | 888 | 2 |
| 56 | 289.340 | 888 | 3 |
| 84 | 64.740 | 888 | 1 |
| 84 | 321.983 | 888 | 2 |
| 84 | 361.557 | 888 | 3 |
| 88 | 213.167 | 888 | 1 |
| 88 | 350.976 | 888 | 2 |
| 90 | 386.325 | 888 | 1 |
| 90 | 376.431 | 888 | 2 |
| 90 | 358.059 | 888 | 3 |
| 98 | 177.878 | 888 | 1 |
| 98 | 302.260 | 888 | 2 |
| 98 | 357.384 | 888 | 3 |
| 112 | 100.208 | 888 | 1 |
| 112 | 167.345 | 888 | 2 |
| 112 | 232.362 | 888 | 3 |
| 140 | 82.671 | 888 | 1 |
| 140 | 151.928 | 888 | 2 |
| 140 | 187.265 | 888 | 3 |
| 168 | 256.655 | 888 | 1 |
| 168 | 120.964 | 888 | 2 |
| 168 | 112.484 | 888 | 3 |

**ANNEX 6: STEADY-STATE CALCULATION FOR THE SIMPLEST TK MODEL**

The following equation (Eq. (7) in the manuscript):

$$\frac{dC_p(t)}{dt} = k_{uw} \times c_w - k_{ee} \times C_p(t) \text{ for } 0 \leq t \leq t_c$$

At the steady state, it can be written as:

$$\frac{dC_p(t)}{dt} = 0 \Leftrightarrow k_{uw} \times c_w - k_{ee} \times C_{p_{ss}} = 0 \Leftrightarrow C_{p_{ss}} = \frac{k_{uw}}{k_{ee}} \times c_w$$

with $C_{p_{ss}}$the concentration of the contaminant at steady state in the organism.
